# Metabolic reprogramming dynamics in tumor spheroids: Insights from a multicellular, multiscale model

**DOI:** 10.1101/485201

**Authors:** Mahua Roy, Stacey D. Finley

## Abstract

Mathematical modeling provides the predictive ability to understand the metabolic reprogramming and complex pathways that mediate cancer cells’ proliferation. We present a mathematical model using a multiscale, multicellular approach to simulate avascular tumor growth, applied to pancreatic cancer. The model spans three distinct spatial and temporal scales. At the extracellular level, reaction diffusion equations describe nutrient concentrations over a span of seconds. At the cellular level, a lattice-based energy driven stochastic approach describes cellular phenomena including adhesion, proliferation, viability and cell state transitions, occurring on the timescale of hours. At the sub-cellular level, we incorporate a detailed kinetic model of intracellular metabolite dynamics on the timescale of minutes, which enables the cells to uptake and excrete metabolites and use the metabolites to generate energy and building blocks for cell growth. This is a particularly novel aspect of the model. Certain defined criteria for the concentrations of intracellular metabolites lead to cancer cell growth, proliferation and necrosis. Overall, we model the evolution of the tumor in both time and space. Starting with a cluster of tumor cells, the model produces an avascular tumor that quantitatively and qualitatively mimics experimental measurements of multicellular tumor spheroids. Through our model simulations, we can investigate the response of individual intracellular species under a metabolic perturbation and investigate how that response contributes to the response of the tumor as a whole. The predicted response of intracellular metabolites under various targeted strategies are difficult to resolve with experimental techniques. Thus, the model can give novel predictions as to the response of the tumor as a whole, identifies potential therapies to impede tumor growth, and predicts the effects of those therapeutic strategies. In particular, the model provides quantitative insight into the dynamic reprogramming of tumor cells at the intracellular level in response to specific metabolic perturbations. Overall, the model is a useful framework to study targeted metabolic strategies for inhibiting tumor growth.

## Introduction

Cancer evolution occurs as an interactive and collaborative process between cellular populations and the extracellular environment spanning over multiple spatial and temporal scales. Understanding the molecular peculiarities of a malignant tissue relative to healthy tissue primarily involves various genome sequencing protocols of a single region of the carcinoma or a single tumor biopsy averaged over the complete tumor mass. However, this type of molecular data fails to capture the regional heterogeneities in a tumor mass. Often, these regional differences within the primary tumor mass mean that portions of the tumor have evolved to be genetically different from the initial cells that were present when tumor growth began (1, 2). Both inter- and intra-tumor heterogeneities influence how the tumor as a whole behaves in different microenvironments, leading to different grades of tumor progression. Hence, it is necessary to understand not only the temporal evolution of the tumor volume but also the spatial heterogeneities within the tumor. Most studies of tumor heterogeneity have focused on the evolution of regional differences within a tumor due to the tumor’s extracellular environment (including nutrient availability and physical characteristics such as matrix stiffness (3)), the mechanical properties of the cancerous cells themselves (4), or the evolution of cellular phenotypes (5, 6). However, few studies investigate the internal metabolic dynamics of individual cells, which might influence how the cells behave in diverse extracellular environments, how they interact with other cells, and how they transition between different cells types (7, 8).

Studying spatial heterogeneity in the context of metabolism is particularly important in pancreatic cancer. Pancreatic adenocarcinoma is one of the most aggressive forms of cancer mostly due to metastatic dissemination and resistance to chemotherapy (9). Pancreatic cancer cells rewire their metabolic network to support the energetic and biosynthetic demands of the exponentially growing tumor. This altered metabolism is a hallmark of cancer and can be linked to resistance to therapies (10). Hence, it becomes important to understand the metabolic dynamics of a pancreatic cancer cell not as homogeneous distribution, but within a heterogeneous tumor volume, where each cell may be subject to nutrient deprivation and has the ability to reprogram their pathways to their own individual benefit.

The behavior of a population of cells is a collective phenomena arising from a complex network of mutually interacting cells, the basic entity of biology, that follows a set of stereotypical or stochastic rules. Cellular signaling operates over several orders of magnitude in a spatio-temporal scale where both extracellular and intracellular dynamics are involved (11, 12). A key challenge is to consider and understand the interconnection of the extracellular signals to the intracellular process and how these processes couple together to prompt cellular response and decisions (13, 14). While most metabolic reactions occur on the order of fractions of seconds to seconds, the end result of phenotypic changes or cellular growth and differentiation takes place on the timescale of hours. Hence, timescale separation is a crucial consideration for dynamic network analysis and spatio-temporal evolution of network systems (15).

Cell-based models are a class of computational models that can mimic the biophysical and molecular interactions between cells. Thus, these models are essential tools to simulate heterogeneous biological behavior. While continuous models are capable of explaining the overall behavior of the tumor as a whole (16), a discrete model (17) can capture the stochasticity of the individual cell’s behavior within the tumor, leading to differential spatio-temporal evolution and inter-tumor heterogeneity. In this type of modeling, each cell’s decisions are simulated based on its current state, local environment, and interactions with its neighbors. Cell-based modeling employs a hybrid approach combining the aspects of discrete cell behavior with a continuum environment, both directly influencing each other. These models are comprised of a set of rules that guide the cell’s decisions and the collective “morphodynamic” behavior of the collection of cells (18). These rules are based on the principles of thermodynamics and mechanics and relevant aspects of physical sciences describing cell-cell interactions and the ensemble of cells. Dynamic cell-based models have been previously used to study how the collective behavior of individual cells drives tissue-level processes, such as in tumor development and angiogenesis (19, 20).

In this study, we use a multiscale cell-based approach to predict how each individual cell’s detailed intracellular metabolic dynamics influence the evolution of the tumor as a whole. We further predict the effects of drugs inhibiting specific enzymatic reactions in inhibiting the growth of the entire tumor. The novelty of our model of avascular tumor growth described here lies in the incorporation of a detailed mechanistic metabolic network (21) within each cell unit. We couple the cells’ behaviors to the diffusion of multiple nutrients, where the cellular growth and death rates are a function of ATP generation rate of a cell unit. The model is able to predict the effects of various therapeutic strategies that target individual cells’ metabolism, i.e., the resulting tumor growth following changes in the expression levels of enzymes (simulated by knockout or knockdown of the enzyme activity), which alter the flux distributions within the intracellular metabolic network, and hence tumor growth. We apply the model to explore cell phenotypic changes and identify which enzyme knockdowns have the maximum influence on reducing tumor growth. Given the intracellular molecular detail of our model, we are able to explain the mechanisms behind each metabolic perturbation and predict their effects at the intracellular level. Our work contributes to a deeper understanding of the effects of metabolic strategies to regulate tumor growth.

## Materials and methods

### Modeling Platform

We model the spatio-temporal early evolution of a multicellular pancreatic tumor spheroid using the Cellular Potts model via an open source simulation environment called CompuCell3D (henceforth referred to as CC3D) (22). Cellular Potts modeling makes it easier to understand both the intra- and inter-tumor heterogeneity due to differences at the cellular level. The modeling environment is based on Glazier-Graner-Hogeweg (GGH) model (23, 24), where each individual cell can interact with each other and its extracellular environment to generate the collective cell behavior.

### Model Overview

Our multiscale model of tumor growth incorporates a spatio-temporal evolution of an avascular tumor spheroid (25) within a two-dimensional stromal tissue compartment (Figure 1). This involves a description of central carbon metabolism within a small, homogeneous cluster of cells (“*generalized cell*”) via a detailed kinetic model. Cells are assumed to be chemically homogeneous internally, i.e., no chemical diffusion is modeled within the intracellular space. The local extracellular concentrations of glucose, glutamine, lactate and oxygen are described with reaction-diffusion equations. The generalized cells are located on a two-dimensional lattice, which represents the tissue domain and the extracellular matrix. Each cell in our model is its own decision-making unit, where decisions are based on the cell’s immediate extracellular conditions, interactions with neighboring cells and its intracellular condition.

**Fig. 1.**
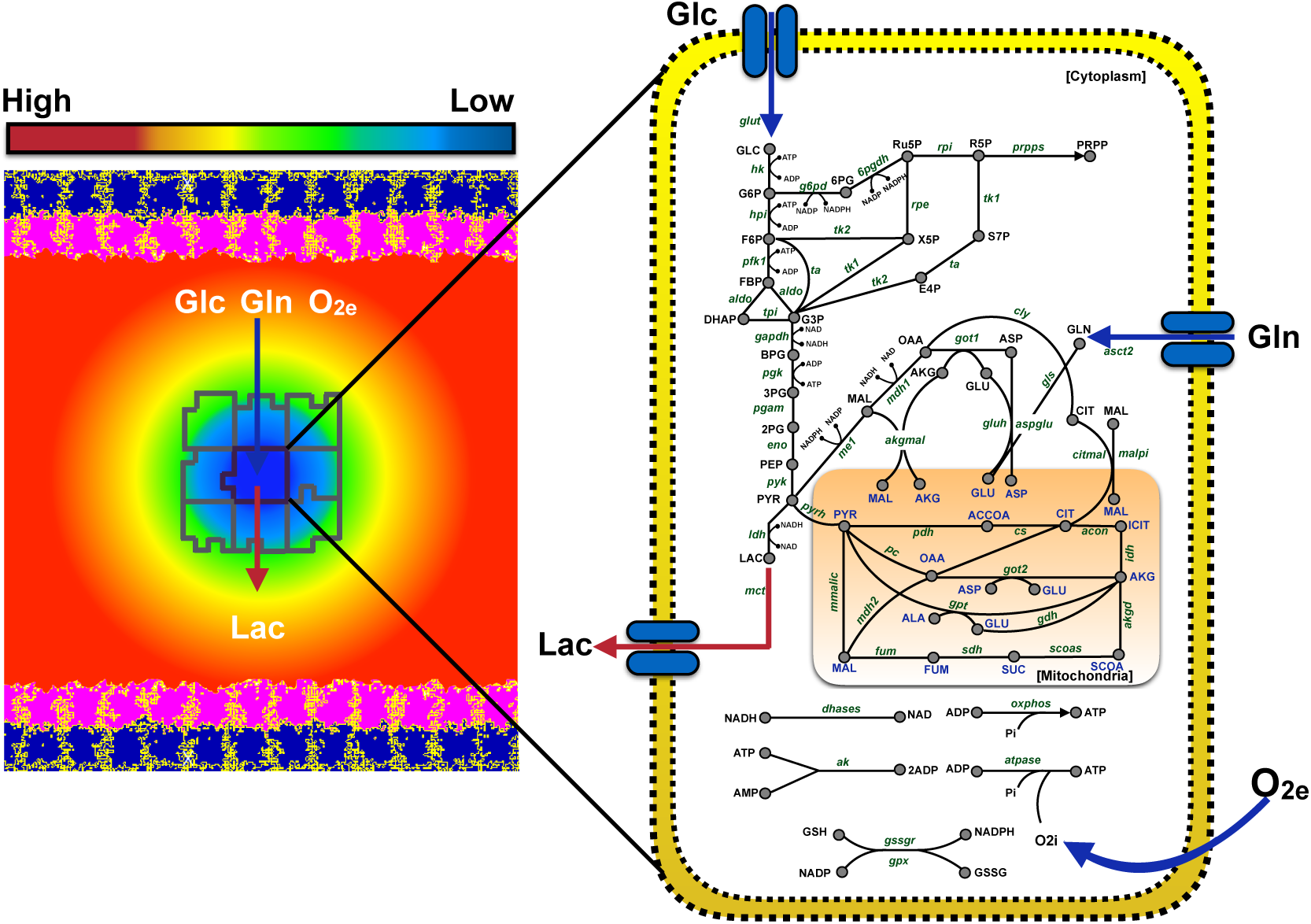
Schematic of the simulated tumor environment: The cellular environment is comprised of a continuous stromal tissue flanked on both sides by an epithelial layer. This epithelial layer is comprised of two cell types - Extracellular Matrix (pink) and Basal (navy blue). The epithelial layer secretes three main nutrients - glucose, glutamine and oxygen. The tumor cells (outlined in grey) are discrete entities that reside within the stromal tissue. Tumor cells take up glucose, glutamine and oxygen, and they secrete lactate into the stromal tissue. The diffusion of these nutrients is modeled using partial differential equations (illustrated by the gradient field, ranging from low concentration (blue) to a high concentration (red)). Each individual tumor cell has a detailed intracellular metabolic network, which takes the extracellular glucose, glutamine and oxygen concentrations as inputs to form ATP with lactate as a by-product of glycolysis. The dynamics of the metabolic network within each cell is modeled using ordinary differential equations. Cell sizes are not to scale.

### Cell Properties

In a Cellular Potts model (implemented using CC3D), each generalized cell has a unique identity number and is a dynamic entity that responds actively to the nutrient gradients in its environment. The cells have adhesive properties to their neighbors, change their shape (volume) and surface properties and exert forces on each other and their environment. They grow and proliferate via mitosis (division) and die due to necrosis when they shrink and disappear. Each cell has a type given by *τ* (*σ*(*x*)) where *σ*(*x*) is the unique identity of the cell at a spatial location, *x*. There can be many cells of the same type, but each has a different identity.

Each generalized cell is associated with an effective energy term or Hamiltonian, **H**, that encapsulates all its cellular properties (volume and surface area), interactions with other cells (adhesion to neighbors), and motion (chemotaxis). The effective energy also includes the set of partial differential equations and boundary conditions that describe the evolution of extracellular nutrients’ gradients in the tumor microenvironment.

### Cell Dynamics

The cell configuration evolves through stochastic changes (Monte Carlo method) at individual sites to minimize effective energy. These are called the *Voxel Copy Attempts*. For the motion of a cell from a source site “*j*” to a target site “*i*”, the change in energy (ΔH) required to copy the index in the source site voxel onto the target site voxel is calculated. The probability of a voxel copy attempt to be accepted or not is given by

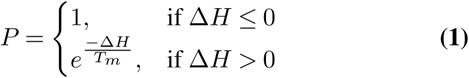

Here, *T_m_* is the temperature, which represents the amplitude of cell membrane fluctuations. The higher the energy of a configuration, the less probable the voxel copy attempt. Overall, the cellular motion (chemotaxis) is driven by a lowering of the effective energy.

Three major factors determine cellular growth and death rates and cell state transitions in the model: (a) local ATP generation due to the intracellular metabolic pathways that utilize glucose, glutamine and oxygen (26), (b) acidity produced by local lactate secretion (27), and (c) growth-induced or residual solid stress within the interior of the tumor (28–31). These factors collectively influence the spatial distribution of proliferating, quiescent, and necrotic cells within the tumor, thus defining the macroscopic tumor morphology. The detailed intracellular kinetics of metabolism in a generalized cell is represented by an ordinary differential equation (ODE) model that relates the cellular uptake of nutrients for the generation of cellular energy (ATP) leading to cell growth or death. The uptake rate of each nutrient is determined by the kinetic parameters of the ODE model as well as the amount available at the center of mass of the generalized cell. While the baseline ODE model framework is based on literature sources (21, 32), additional features such as tumor response to acidosis and compressive stress (33) were added. The various components of the model, and the parameters associated with them, are as described below.

### Grid Space and Cell Types

The spatial extent of the simulations involves a lattice grid with dimensions: 400 × 400 × 1 pixels. Voxels on this fixed cell lattice represent generalized cells. The correspondence of pixel units of a generalized cell to length and volume units are given in **Table S1 in S1 Text**. Based on the volume of a generalized cell, a universal scaling factor is imposed for all parameters of the model which, assumes that 5 mM corresponds to 0.32 fmol/voxel (32).

There are a total of eight different cell types in the tumor environment. The lattice space is occupied by the **Medium** cells, which represent a stromal compartment. While the **Medium** cells are continuous, the other cell types are discrete and represent spatially extended domains on the space lattice. The stromal compartment is bound on each end by an epithelial layer, occupying 40 pixels of space. The epithelial layer is comprised of two cell types: the lower layer is the extracellular matrix (**ECM**) cells, and the upper layer is **Basal** cells. While lactate is only secreted by the stromal compartment, the epithelial layer and the stromal compartment both secrete glucose, glutamine and oxygen. The schematic of the model is shown in Figure 1.

The model assumes that the growth of a malignant tumor is initiated from a small cluster of destabilized and disordered cells, which can be either quiescent or possess proliferative capacity, based on the availability of the nutrients. These quiescent and proliferating tumor stem cells originate at the center of the lattice with a periodic boundary condition imposed on both *x* and *y* directions. As time progresses, the tumor can evolve to include five different types of cells - **PCancer** and **PStem** are the proliferating tumor and stem cells, respectively; **QCancer** and **QStem** are the quiescent tumor and stem cells, respectively; and **Necrotic** are the cells on the verge of cell death (apoptosis or necrosis). Necrotic cell types at the periphery of the tumor eventually shrink in volume and disappear, their space taken by other cell types. This mimics the phenomenon of apoptosis, where the cell disintegrates with the rupture of the cell membrane due to the action of immune cells. The necrotic cells at the core of the tumor shrink in size to one fourth of their original volume; however, they do not disintegrate, due to spatial constraints preventing immune cells from accessing the inner region of the tumor to clear the cell’s debris. In both cases of transitioning into a Necrotic cell, the cell transfers its lactate to the microenvironment, increasing the local acidity.

### Nutrient Field and Chemotaxis

The cells grow in response to availability of nutrients (glucose, glutamine, oxygen and lactate) in the extracellular space. These concentrations of the extracellular nutrients are influenced by the cells’ secretion, uptake and chemotaxis. The nutrient field is initiated with the assumption that there is no consumption by the tumor cells. The stromal cells and the epithelial layer in the model secrete nutrients at a constant rate and have a constant consumption rate such that the physiologically normal ranges of these nutrients are maintained at all times. We assume a constant secretion rate of nutrients at an arbitrary value of *α*. The consumption rate, *ε*, is calculated for each nutrient using the same approach as Swat *et al.*(32), so as to maintain the field at a steady state concentration (*G*(*x*)) in the absence of tumor cells.

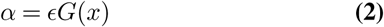

The normal physiological concentrations of the nutrients in blood and their corresponding diffusion constants, along with the corresponding consumption rates, are tabulated in **Table S1 in S1 Text**. The nutrient diffusion within the lattice is modeled using the `SteadyStateDiffusionSolver` within CC3D. This solver implements the Helmholtz equation as a time-independent form to solve the diffusion of the nutrients that have very high diffusion constants.

In the model, a cell can uptake all three nutrients (glucose, glutamine and oxygen), when available, at its center of mass. This nutrient uptake drives the cell’s intracellular metabolism. When glucose is no longer available (*Glu_out_* `= 0`), lactate is consumed as an alternative source of carbon (27). At a normal glucose concentration, the lactate uptake is null. The nutrients’ field gradient is a net result of secretion and decay by the epithelial layer and consumption by the cells.

The field gradient causes the proliferating cells to chemotax, as glucose, glutamate and oxygen are chemo-attractors. However, there is no preferential chemotaxis towards a specific nutrient, as all three nutrients have the same chemotaxis strength coefficient, *λ*. The contribution to the cell’s effective energy, *H*, due to chemotaxis is given by

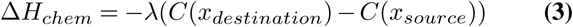

Chemotaxis is simulated by voxel copy attempts, similar to the stochastic fluctuations in the cell configuration (see above). Here, if the nutrient concentration at the destination pixel is more than the concentration at the source pixel, the voxel copy attempt is accepted, as it leads to a lowering of energy. This in turn leads the cell movement towards the chemo-attractant. If the nutrient concentration at the destination pixel is lower than that at the source pixel, the probability of the voxel copy being made follows equation (1) shown above.

### Cell Volume and Cell Surface

Each cell has a defined volume (V) and a deformable shape and surface area (S), which contribute to the effective energy, *H*, as follows:

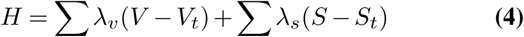

An energy penalty increases with the deviation of the cell’s instantaneous volume and surface area from a designated target volume (*V_t_*) or target surface (*S_t_*), respectively, which imposes volume and surface area constraints on the cells. The *λ*_*v*_ and *λ*_*s*_ parameters can be correlated to the Young’s modulus, with higher values reducing the magnitude of fluctuations of a cell’s volume or surface area about its target value. The *λ* values chosen for tumor cell types (PCancer, PStem, QCancer, QStem and Necrotic) are given in **Table S2 in S1 Text**.

### Interaction with Other Cell Types

The contact adhesion or repulsion of a cell with its nearest neighbors contributes to the effective energy, *H*, via the function

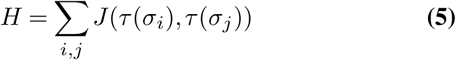

Higher or more positive value of the coefficient *J* leads to repulsive behavior among cell types, whereas more negative or lower *J* values result in greater adhesion. Hence, the cells that adhere together should have a low *J* value in order to lower their overall energy. Each cell type has different grades of adhesion with identical or different cells, as given in **Table S3 in S1 Text**. These values are chosen relative to each cell type. For example, to limit necrosis to the core of the tumor, the adhesion coefficient of the necrotic cells with stromal and epithelial cells is chosen to be high relative to the other cell types.

Altogether, the net effective energy for the generalized tumor cells due to cell properties, interactions with other cells, and chemotaxis is given by

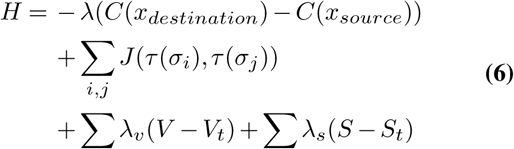

### Stability of the Epithelial Layer

The epithelial layer is constrained with higher target volume and surface area values, as given in **Table S2 in S1 Text**. The layers of ECM and Basal cells are constrained using the *FocalPointPlasticity*, a length constraint imposed to maintain the maximum number of cell-cell junctions. The energy contribution to the effective energy, *H*, is given as

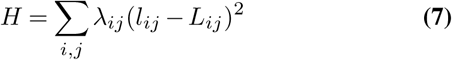

where *l_ij_* is the distance between the center of masses of the two neighboring cells, and *L_ij_* is the target distance between them. The strength of the links between the cells is given by the coefficient *λ*_*ij*_. An energy penalty is incurred when the distance between the center of masses of the cells reaches *MaxDistance* and the link breaks. However, cells in the process of reconnecting can also come closer to other separated neighboring cells and restore the broken link in order to lower the energy. The parameter values governing this stability of the epithelial layers are tabulated in **Table S2 in S1 Text**. The total effective energy, *H*, for the epithelial layer is therefore given by

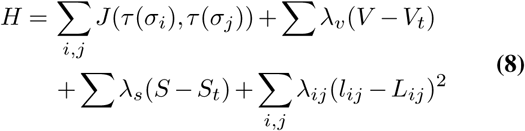

The complete set of parameters mentioned above remain fixed and are tabulated in **Table S2 in S1 Text**.

### Simulation Time

Due to the discrete and multiscale aspects of this modeling approach, it is computationally very expensive and infeasible to simulate the growth of large tumors. Hence, we limited the model simulations to mimic the growth of an avascular tumor, focused on reproducing the volume fold change as observed in multicellular tumor spheroids. Tumor evolution is simulated for a total time period of 25 days, based on the experimental data for multicellular tumor spheroid growth of various cancer cell lines (34–36). In CC3D, the generalized cells migrate at the speed of 0.1 pixel per Monte Carlo Step (MCS) unit. Based on the tumor cell migration speed of 4 *µ*m/hr observed experimentally(37), the correspondence of real time to MCS is 6 minutes. This brings the total simulation time to be 6000 MCS, equivalent to 25 days.

Cellular decisions and interactions take place on different time scales, quite different from a cell’s intracellular dynamics. While the metabolic reactions are fast, and metabolites in the network achieve steady state in several minutes, the growth of the tumor is on the order of days. Hence, we assumed that the cellular decision-making processes, which are mediated by the intracellular dynamics, occur on an intermediate time scale of hours(38). Thus, the “fast” reactions (the intracellular metabolic dynamics) could be considered to be at quasi-steady, relative to cellular decisions and tumor growth. We integrate the different time scales by updating the various processes at different times: the intracellular dynamic concentrations are updated every MCS (every 6 minutes), and the cellular decisions of growth, cell state transition and mitosis occur every 5 hours. We describe these processes below.

### Intracellular Dynamics

Both proliferating (PCancer and PStem) and quiescent cells (QCancer and QStem) have an elaborate intracellular metabolic network, simulated though an ODE model (21), implemented as SBML code. This ODE model initiates with the uptake of glucose, glutamine, oxygen and in certain specific cases (as mentioned above), lactate from the stromal compartment (Medium cells). As a result of metabolizing those nutrients, the ODE model produces ATP, which is then recorded for each cell (Figure 1). These simulations are continuous. That is, after each MCS, the cell’s intracellular state is updated with its final state from the previous MCS. This updating occurs every 6 minutes.

### Cell Growth, Necrosis and Mitosis

Cell growth in the model is limited to proliferating cells (PCancer and PStem), with their growth dependent on the amount of ATP produced by the cell (21). The rate of production of ATP depends on the total amount of intracellular glucose and glutamine, as well as additional intracellular components generated from glycolysis, the tricarboxylic acid cycle (TCA) and the pentose phosphate pathway (PPP). Cell growth occurs for a particular cell when the total intracellular concentration of ATP, glucose and glutamine produced by that cell is above a certain growth threshold: *PGrThr* and *SGrThr* for PCancer and PStem cells, respectively. For the cells that meet this threshold, the cell’s target volume is incremented by a constant rate at the time interval for cell growth (5 hours). The new cell volume (*V_tf_*) is given by:

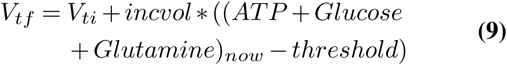

where *V_ti_* is the initial volume, *incvol* is the growth rate in units of *voxels*^2^*/fmol* and *threshold* is *PGrThr* or *SGrThr* for PCancer and PStem cells, respectively.

Cells can become necrotic due to very low production of ATP. Here, we assume that cell necrosis is only a function of the cell’s ATP production. The volume of necrotic cells decreases at a constant rate, *decvol* (given in units of *fmol/voxel*), mimicking cellular apoptosis. The new cell volume is given by:

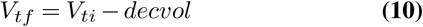

The volume of necrotic cells that appear at the tumor periphery can decrease to zero, as these cells get cleared from the environment or turn into debris. When a necrotic cell develops in the core of the tumor, the cell shrinks following the equation above, but its volume only decreases to one fourth of the size of a proliferating or quiescent cell. In both cases, all of the necrotic cell’s intracellular lactate is transferred from the interior of the cell to the microenvironment, mimicking the rupturing of the cell membrane and increasing the acidity of the local microenvironment.

Cell mitosis (cell division) occurs when a cell’s volume increases such that it reaches the maximum allowable size: *Pvolmaxmit* or *Svolmaxmit* for PCancer or PStem cells, respectively. We use the same values for *Pvolmaxmit* and *Svolmaxmit*, equal to twice the initial cell volume (2.0\**V_o_*).

### Cell Attributes

The cell’s transition from one state to other is dependent on its local environment and intracellular dynamics. We define the **Health** attribute to characterize the health of a cell, which depends on its rate of production of ATP. A cell can also accumulate damage through the **Starvation** and **Acidity** attributes. Starvation is a function of the ATP production and occurs due to low glucose and glutamine. Acidity occurs due to over-exposure to lactate. Finally, a cell accumulates **Stress** due to pressure exerted by nearby cells over a prolonged period of time. Starvation, Health and Acidity acquired by the cell are calculated via a fitness factor, *F*, as follows:

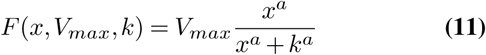

where *x* is ATP concentration (for Health and Starvation) or lactate concentration (for Acidity). *V_max_* is the maximum ATP or lactate production by the cell. For a quiescent cell, the *V_max_* is 75% of that of a proliferating cell, *a* is the Hill coefficient, and *k* is the Michaelis constant. For the calculation of Health, *F* is multiplied by a coefficient, `exp(C)`, a measure of utilization of ATP and intracellular glucose and glutamine for cell growth. At each time step, the cell attributes of Starvation, Health, and Acidity are incremented by an amount such that:

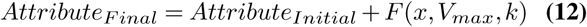

In the case of Stress, the cell attribute is increased linearly by an amount equal to *StressIncrement*. Since the amount of stress increase is dependent on the probability of a cell having a specific number of neighbors, we assigned a linear increment each time the event occurs, instead of an increasing Michaelis Menten function. After each state transition, the cell’s attributes are reinitiated to zero.

### Cell State Transitions due to ATP Production

The cell’s production of ATP due to metabolizing glucose, glutamine and lactate for a specified period of time enables the transition from quiescent to proliferating. Specifically, the thresholds to acquire Health are calculated based on the cell’s consistent production of ATP, along with intracellular glucose and glutamine concentrations over a time period (*Total_time_*).

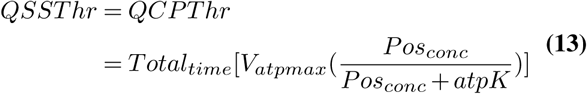

where *QCPThr* and *QSSThr* are the threshold values for the conversion of QCancer and QStem cells to proliferating cells, PCancer and PStem cells, respectively. *V_atpmax_* and *atpK* are the rate constants needed such that the threshold value saturates with increasing *Pos_conc_*, following Michaelis-Menten kinetics. *Pos_conc_* is the sum of the concentration of glucose, glutamine and lactate.

### Cell State Transitions due to Lack of ATP

The cells start to acquire Starvation when ATP production is lower than a certain threshold (*atpD*) and otherwise acquire Health. The threshold value for a proliferating cell to become necrotic, *PNeThr*, is calculated based on the cell’s exposure to very low ATP (*Neg_concAT P_*) persistently for a time period (*Total_time_*). When the proliferating cell crosses this threshold, it transitions into a necrotic state, an irreversible change. The threshold value is calculated as:

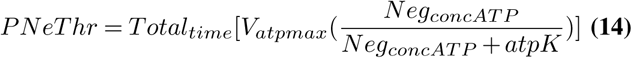

where *V_atpmax_* and *atpK* are the rate constants needed such that the threshold value saturates with increasing *Neg_concAT P_*, following Michaelis-Menten kinetics. *Neg_concAT P_* is a fixed concentration of ATP.

We assume that the quiescent and stem cells take a longer time to transition into the necrotic state, as compared to proliferating cells, and hence have higher thresholds, as tabulated in **Table S2 in S1 Text**.

### Cell State Transition due to Acidosis

The microenvironment invariably becomes acidic in nature due to an increased lactate secretion by necrotic cells as the tumor grows. This acidic environment, in turn, leads to more necrosis if cells experience an external lactate concentration greater than a certain maximum threshold. Both proliferating and quiescent cells become necrotic from acidosis, i.e., continuous overexposure to excess lactate in the microenvironment. The threshold values for acidosis are calculated in a similar way as those for the lack of ATP, with the assumption that the cells are in a continuous exposure of lactate, *Pos_concLac_*, greater than a threshold, *LacDeath* and this exposure persists for a definite time period, *Total_timelac_*. As above, it is assumed that the threshold for acidosis to set in for quiescent and stem cells is higher than that of proliferating cells.

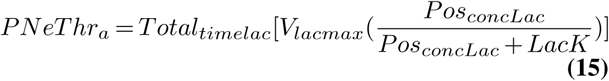

where *V_lacmax_* and *LacK* are the rate constants needed such that the threshold value saturates with increasing *Pos_conc_*, following Michaelis-Menten kinetics. The Michaelis constant *LacK* has a constant value 1 mM (0.064 fmol/voxel).

### Cell State Transition due to Compressive Stress

The cells at the core of the tumor transition into necrotic cells due to stress induced by the neighboring cells. The cell’s Stress factor is an added attribute that leads to necrosis and starts developing at the core of the tumor, when the cells consistently have more than a certain number of surrounding neighbors, *N*. Every time step that a cell has more than *N* neighbors, *StressIncrement* is added to the cell’s Stress attribute. When the cell’s Stress crosses a threshold, *StressThr*, it becomes necrotic.

### Cell Mitosis: Senescence and Differentiation

Cell mitosis is a function of cellular volume. When a fully grown, proliferating cancer or stem cell reaches its doubling volume, it divides. As this cell (referred to as the “*parent cell*”) divides, its voxels are equally divided between the parent cell and daughter cell, and both cells acquire quiescence. The daughter cell inherits all properties and the last intracellular metabolic state of the parent cell. The cell attributes (Starvation, Acidity, Stress and Health) are reinitiated to zero for both the parent and daughter cells.

Upon division, both the proliferating parent cell and its daughter cell become quiescent cancer cells. Proliferating stem (PStem) cells follow a different process. When a PStem cell divides, this parent cell becomes a quiescent stem (QStem) cell. However, its daughter cell has a probability of being a quiescent stem (QStem) or quiescent cancer (QCancer) cell. This is decided by a probability, *probstem*. At each division step, a random number chosen between 0 and 1 is compared with the probability of transitioning into a Stem cell, *probstem*. If the random number is less than or equal to *probstem*, the daughter cell becomes a quiescent stem cell, otherwise it becomes a quiescent cancer cell.

For a proliferating cancer cell, mitosis is arrested (*i.e.*, senescence occurs) when the cell has undergone a maximum number of divisions(39, 40), after which the cell becomes necrotic. We assume proliferating stem cells also have a maximum number of divisions. To keep an account of when senescence should occur, a counter, *temp*, is introduced. The value of *temp* for each proliferating cell is chosen as a random number from a Gaussian distribution with a mean of *maxdiv* and standard deviation of two. When a proliferating cell reaches its doubling volume, but that cell has already undergone a number of cell divisions exceeding the value of *temp*, the cell does not divide and instead becomes necrotic.

The complete flowchart depicting all of the cell state transitions described above is shown in Figure 2.

**Fig. 2.**
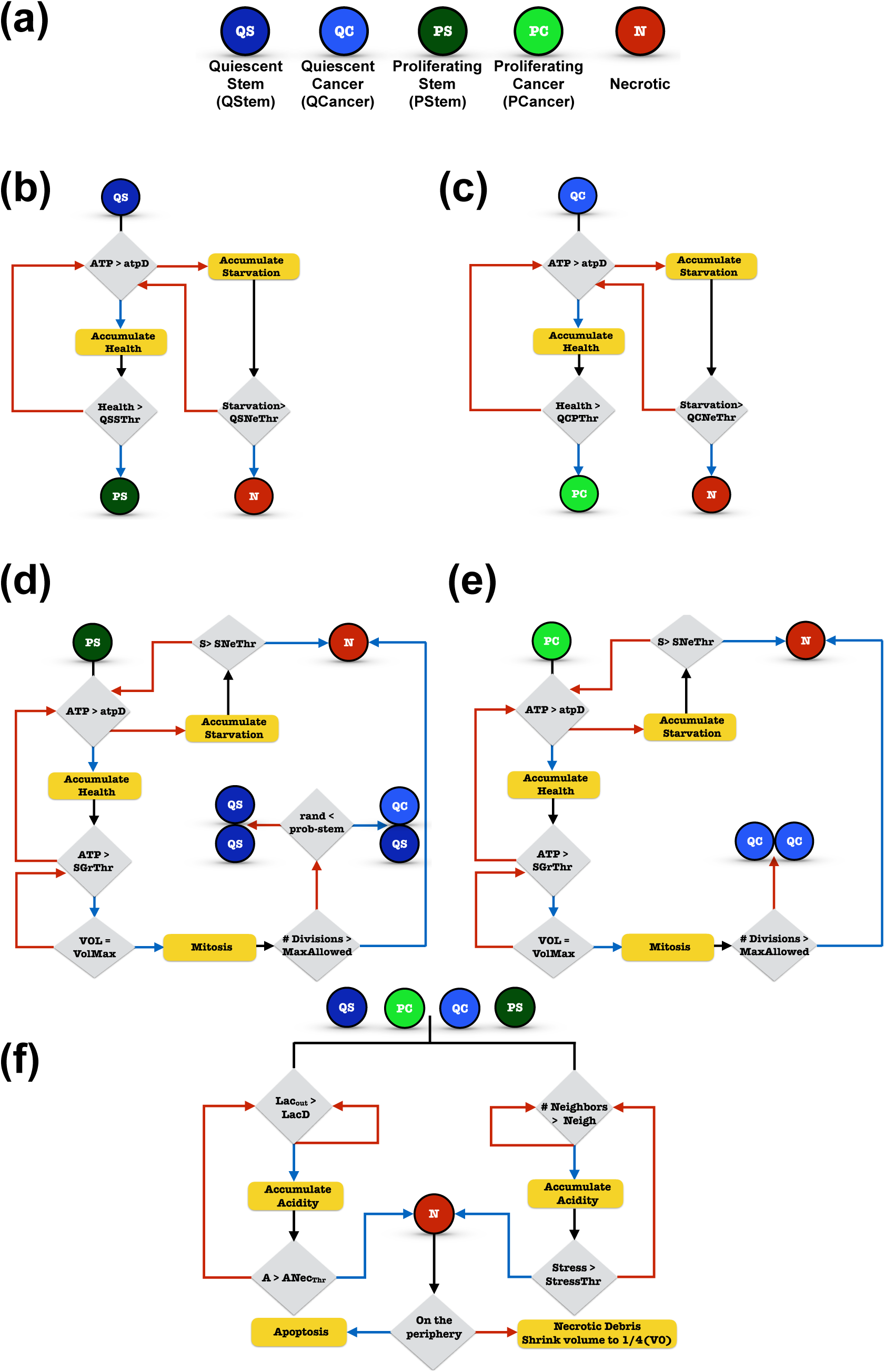
Cell Types and Cell Transitions: (a) Discrete cell types in the model. Rules for the transition of (b) a quiescent stem cell, (c) a quiescent cancer cell, (d) a proliferating stem cell or (e) a proliferating cancer cell. (f) Rules for the transition of all four viable cell types into a necrotic cell.

### Model Calibration through Parameter Estimation

The results from numerical simulation of a mathematical model are dependent on the values of the model parameters. Hence, it becomes important to determine the parameter values of a mathematical model to ensure the simulated results recapitulate experimental observations. In this work, the model is trained and calibrated to match the experimental data of multicellular tumor spheroid growth (34–36), shown in Figure S1. Due to the computational expense, it was not feasible to start with the actual number of cells present within a cell culture (approximately 105) or to simulate the same number of generalized cells as found in a real tumor. With the available computational resources, we instead focused on capturing the experimentally-measured fold-change in tumor spheroid volume. We collect data for the growth of multicellular tumor spheroids for various cancer cell lines. The model parameters were sampled within the ranges tabulated in Table 1 and trained so as to reproduce, both quantitatively and qualitatively, the growth profile of a multicellular spheroid. Our intracellular metabolic core model is trained to data from PaTu8988T cells (21); however, we were unable to find tumor spheroid growth data for this specific cell line. Thus, we chose a 10-fold change in tumor volume in approximately 15 days to be the standard for the model-simulated tumor growth.

**Table 1.** Range over which the parameters are varied in iterations to calibrate the model for multicellular tumor spheroid growth: Latin hypercube sampling is used to generate 100 sets of parameter values within the specified minimum an maximum values. The model was simulated 100 times with different combinations of parameter values. The ideal parameter set generates a 10-fold change in tumor volume in approximately 15 days (“baseline” parameter values, last column).

| Parameters | Minimum | Maximum | Optimized set of parameters |
| --- | --- | --- | --- |
| incvol | 10 | 50 | 15.0 |
| decvol | 0.001 | 0.1 | 0.025 |
| PGrThr | 0.0032 | 0.192 | 0.09983 |
| SGrThr | 0.0032 | 0.192 | 0.09983 |
| StressIncrement | 0.25 | 1 | 0.95 |
| StressThr | 20 | 100 | 50.5 |
| N | 7 | 10 | 6 |
| $V_{atpmax}$ | 1.68e-03 | 4.2 | 0.6 |
| atpD | 0.064 | 0.032 | 0.021 |
| $Total_{time}$ | 50 | 240 | 95.0 |
| $Neg_{concATP}$ | 0.0001 | 0.015 | 0.00095 |
| $Pos_{conc}$ | 0.0064 | 0.192 | 0.0628 |
| $V_{lacmax}$ | 1.11e-02 | 27.75 | 15.393 |
| LacDeath | 0.1256 | 2.75 | 1.68 |
| $Pos_{concLac}$ | 0.1256 | 2.75 | 2.22 |
| $Total_{timelac}$ | 50 | 240 | 168.2 |
| maxdiv | 4 | 10 | 9 |
| probstem | 0.1 | 0.8 | 0.5 |
| C | 6 | 11 | 6 |

We varied several model parameters in order to match the 10-fold change in the volume of the tumor spheroids. Specifically, model parameters that could not be based on a literature source, were adjusted within a given range to reproduce the experimental fold-change. These model parameters were categorized as internal kinetic parameters, concentration threshold parameters and external parameters that depend on the tumor microenvironment. An initial range for each parameter was defined (Table 1) based on literature values (21, 32), and the parameter space was sampled using Latin hypercube sampling (LHS). Multiple initial runs were conducted to identify the correct set of parameters using LHS within the parameter space defined by the range given in Table 1. Out of 100 sets of parameters, we selected the best parameter set, one that produces a tumor that satisfies the following physiologically reasonable criteria: (1) the expected tumor volume (normalized to day zero) exhibits a 10-fold change in approximately 15 days and (2) the tumor exhibits a morphology with significant necrosis (41–44) and an appropriate distribution of the various cell types (proliferating, quiescent and necrotic) (45). Moreover, the actual parameter values also satisfy the following criteria, based on the cell behavior and state transitions described above: (1) the value of *Neg_concAT P_* has to be lower than *atpD*, (2) the value of *Pos_conc_* has to be greater than *atpD* and (3) the value of *Pos_concLac_* has to be higher than *LacDeath*.

The set of parameter values that meet all of these criteria are tabulated in Table 1 and are referred to as the “baseline” parameters throughout the text.

### Results

Cancer initiates at the sub-cellular level through events that involve multiple intracellular components interacting with each other. These interactions regulate cellular decisions of growth, death or cell-state transitions. Our model relies on a robust intracellular metabolism network at the sub-cellular level with metabolic rate parameters trained to pancreatic cancer data (21). The parameters corresponding to cell decisions of proliferation and apoptosis were subsequently trained to match the growth curves of tumor spheroids (see Methods). The simulated growth of the tumor is in good agreement, both quantitatively and qualitatively, with the experimental measurements (**Figure S1 in S1 Text**).

### Model Exploration

#### Tumor Growth and Morphology due to Initial Heterogeneity

In order to get a better idea of the tumor morphology under different conditions, model simulations were conducted iteratively to observe the variance in the tumor growth profile with different initial configurations of the generalized cells. We started with five different cases representing different amounts of initial tumor heterogeneity, depending on the number of cell types present in the initial cluster (Figure 3(a)). Case 1 represents the most heterogeneous scenario with all four cell types present (PCancer, PStem, QCancer and QStem). In Case II, only proliferating cells (PCancer and PStem) are initially present. Case III includes only quiescent cells (QCancer and QStem). In Case IV, only proliferating cancer cells (PCancer) are present. Finally, for Case V, only proliferating stem cells (PStem) are present. Two other possible cases initiating with only quiescent stem or quiescent cancer cells were not simulated, as we do not expect them to be much different than Case III, which had a combination of both. The trained cellular parameters produce very similar growth curves in all cases (Figure 3(b)), further providing confidence in the calibrated parameters. However, looking at the different cell numbers and their variance with time (Figure 3(c)) is where we observe differences between the five cases. The fraction of each of the different cell types varies for the five cases, particularly during the initial days of growth. However, the large variations in the distribution of the cell types decreases as the tumor grows with time (see also Figure S2 in S1 Text).

**Fig. 3.**
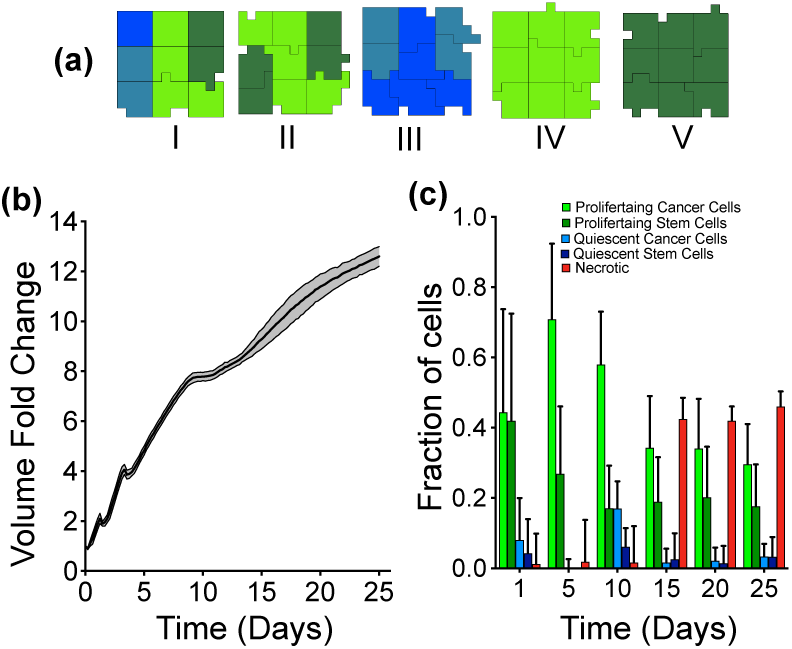
Model Calibration: (a) Five scenarios of initial tumor cell clusters were simulated with different combinations of cells present: (I) All four types of cells (PCancer, QCancer, PStem and QStem), (II) Only proliferating cell (PCancer and PStem), (III) Only quiescent cells (QCancer and QStem), (IV) Only proliferating cancer cell (PCancer) and (V) Only proliferating stem cells (PStem). The ratio between each cell type is randomly generated by CC3D to have a total of nine cells. (b) Model-simulated tumor growth profiles using the optimized parameter set, initiating with the five different cell cluster configurations. Ten iterations for each case were simulated. The black solid line is the mean of all 50 simulations, and the shaded grey area represents the standard deviation of the simulations. (c) Fraction of each type of cell as the tumor grows with time. The bar height is the mean value of all 50 simulations while the standard deviation is represented by the error bars. Time evolution of the number of each cell type differs for five starting cell clusters, shown in Figure S2 in S1 Text.

The development of necrosis has been shown to be an important feature of tumor spheroids. Our model simulations reproduce this feature, where the number of necrotic cells increases with time. Altogether, these results indicate that our model simulations are reliable and that the optimized parameter set is able to successfully reproduce the experimental data, both quantitatively and qualitatively. A representative simulation is provided in S1 Movie.

#### Parameter Sensitivity

Since the cellular parameters are specific to the model, we sought to understand how varying the model parameters affected the growth of the overall tumor. We focused on the intracellular threshold levels of ATP and lactate, which directly influence the cellular behavior and hence, inter-tumor heterogeneity. To do so, we varied two important parameters related to cell death and growth: *atpD*, the ATP threshold value (which influences cell proliferation, cell state transition and cell death) and *LacD*, the lactate threshold (which influences cell death). Since all cellular decisions such as cell growth and death and cell state transitions, depend predominantly on these two parameters, they were varied two orders of magnitude from their baseline values, and the fold-change in the tumor volume was recorded after 25 days of growth. The ATP threshold value (*atpD*) was found to have a more dominant effect in changing the tumor volume. This parameter determines when quiescent cells transition into proliferating cells. The model predicts that the fold-change in tumor volume is two times higher than the calibrated volume (baseline) as the ATP threshold value was decreased by 10-fold from the baseline (Figure 4(a)). The snapshots of tumor mass at the end of 25 days clearly illustrate the increase in tumor volume due to lowering of the ATP threshold value, which leads to more cell growth and division (Figure 4(b)). The threshold value for lactate (*lacD*) is not as influential in affecting a change in tumor volume. This result illustrates that within the simulation time window (25 days), the extracellular concentration of lactate does not reach the limit of inflicting acidosis and subsequent necrosis (found by increasing *lacD*), nor are the cells directly dependent on lactate as a nutrient source (27) (found by decreasing *lacD*). Overall, these simulation results indicate that the growth or shrinkage of the tumor is highly dependent on the amount of intracellular ATP being generated by the cell.

**Fig. 4.**
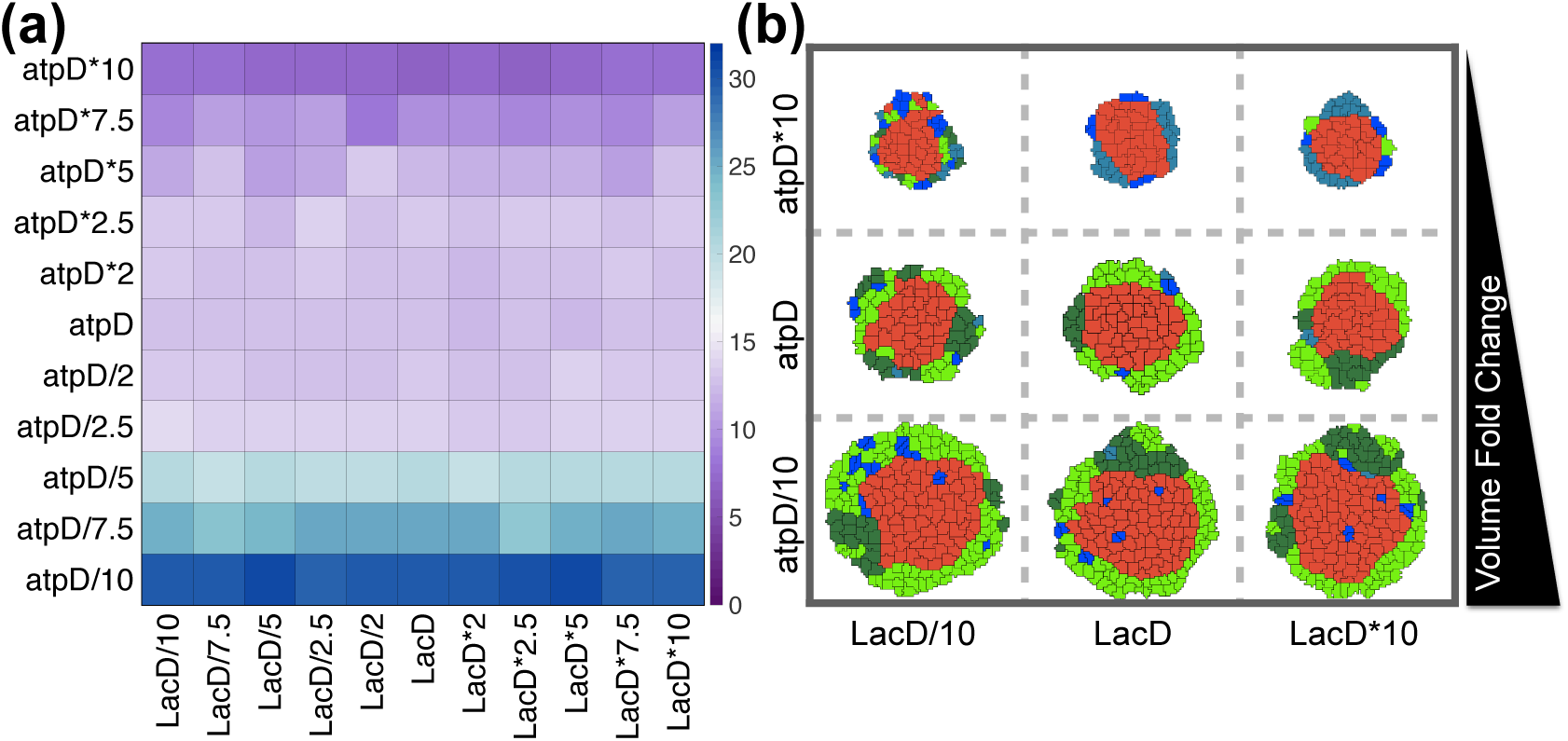
Parameter Sensitivity: (a)The fold-change in tumor volume recorded after 25 days with different values of the threshold parameters *atpD* and *lacD*. For each combination of threshold values, the model was simulated five times, and the mean value is reported in the heat map. (b) Representative snapshots of tumor volume on the 25th day with extreme values of *atpD* and *lacD*, compared to the base value.

### Varying the Initial Metabolite Concentrations

In the context of metabolism, the network of metabolic reactions occurring inside of a cancer cell defines its “metabolic phenotype” (46, 47) that would ultimately lead to inter-tumor heterogeneities within tumors of same origin. Additionally, there is intrinsic cell-to-cell heterogeneity with respect to the initial concentrations of the cells and the activity of the proteins for a population cells. Overall, heterogeneity can result in very different growth patterns among tumors of similar origin, as well as influence their response to therapeutic agents (48). Understanding this heterogeneity is a critical step towards designing improved therapeutic strategies. Thus, we applied the model to investigate the effect of cell-to-cell heterogeneity at the level of intracellular metabolite concentrations.

Our model can successfully capture the variation arising from different initial conditions within the population. We simulated a setting where each cell within the initial cluster of cells starts with a different set of intracellular concentrations. Since the base values for initial intracellular concentrations of metabolites was already validated for a PDAC cell line (21), a tight range of deviation from the base value was maintained. To execute this, we allowed the initial condition of each metabolite to vary around its base value and generated a set of 90 initial concentrations drawn from Gaussian distribution with standard deviation of 0.1. In total, the model is simulated 10 times, where each time, the nine cells start with a slightly different initial intracellular condition. For each simulation run, nine intracellular conditions are randomly chosen, such that each cell within the initial cluster of nine cells has a different initial condition. Even though the initial growth phase of the tumor in these 10 cases are not distinctly different, the variation in tumor evolution increases progressively with time (Figure S3 in S1 Text, blue boxes). Our results highlight the fact that even a modest variation in initial intracellular conditions within a small population of cells can translate into significant differences in the size of the tumor as time progresses. Such heterogeneity can also influence the response to metabolic perturbations, which are investigated in the next section.

### Metabolic Perturbations

In order to identify ideal metabolic targets that can be modulated to reduce tumor growth, we subjected our model to various *in silico* metabolic perturbations. We specifically focused on predicting the effect on tumor volume in response to inhibition of specific metabolic reactions. The use of this kind of computational approach to explore the response of cancer cells could help identify and validate metabolic targets for anticancer therapeutics (49), complementing experimental and pre-clinical studies. Genetic perturbations have been implemented experimentally via complete and partial gene knockdowns (50). Therefore, we enforced the same type of perturbations in our model by altering the reaction velocity parameter (*V_f_*) in the intracellular ODE model, as was done in our previous work (21). This strategy mimics targeted inhibition of a particular enzymatic reactions, for example using shRNA.

Up-regulation of glycolysis is a hallmark of solid tumors (51), where glygolysis is preferred over oxidative phosphorylation to generate ATP for cell growth. In addition, pancreatic cancer cells have been reported to depend heavily on glutamine (52) as an important source of nitrogen and to replenish carbons for TCA cycle intermediates. Hence, we simulated targeted inhibition of each of these pathways. We targeted the reactions OXPHOS (responsible for mitochondrial production of ATP) and ASCT2 (glutamine uptake reaction) to directly inhibit the production of ATP and uptake of glutamine. However, we did not target GLUT1, which would simulate the direct inhibition of glucose uptake. In clinical trials, GLUT1 inhibitors were found to have a limited effect (53, 54), ultimately leading to increased glutaminolysis or increased production of ATP by oxidative phosphorylation. Thus, we chose instead to limit the activity of an identified potential target in the glycolytic cycle, GAPDH (21, 55) (responsible for the conversion of glyceraldehyde 3-phosphate to 1,3-bisphosphoglyceric acid).

We implemented both complete inhibition (at the 0th day of tumor growth) and partial time-based inhibition (0th day, 5th day, 10th day or 20th day) of the three reactions (GAPDH, OXPHOS and ASCT2) independently. During complete inhibition, the reaction velocity *V_f_* of the targeted reaction was given a value of zero. Since the simulations are stochastic, we repeated this complete inhibition 30 times to get a sense for the variability in the model predictions. Partial knockdown or inhibition was implemented by allowing the value of *V_f_* to vary around 0.1 percent of the original value, following a Gaussian distribution with standard deviation of 20%. A total of 30 values were generated from this distribution, and these values were used to repeat the model simulations 30 times, with one value from the distribution used per simulation. We implemented this variability in the partial knockdown to account for the fact that, experimentally, the effect of shRNA is not exact. The recorded fold-change in tumor volume was compared to a control case with no inhibition.

#### Metabolic Perturbation I - Complete inhibition

Complete inhibition of these reactions at the very initial phase of growth has differential effects on the tumor volume (Figure 5). The tumor does not exhibit substantial growth when OXPHOS or ASCT2 are inhibited, similar to experimental results (56). Inhibiting the GAPDH reaction has the initial effect of reducing the tumor growth rate and flattening the growth curve, compared to the control case. However, this effect appears to lessen as time progresses, since the growth curve goes up again between days 20 and 25, matching the slope of the control case during that time period. This is indicative of the cell’s strategy to recover from the growth inhibition as time progresses by exploiting alternative routes for cell growth besides glycolysis.

**Fig. 5.**
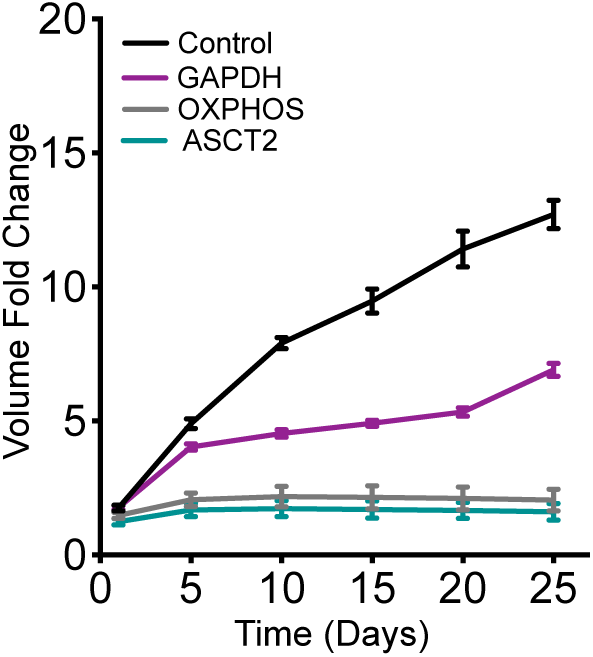
Metabolic Perturbation I - Complete Inhibition: Fold-change in tumor volume with complete inhibition of GAPDH (purple), OXPHOS reaction (grey) and ASCT2 (teal) compared to a control case (black). The error bars represent the standard deviation between 30 iterations of model simulations for each case.

To dig deeper into the causes of this response, we looked at the predicted intracellular concentrations of the metabolites. The detailed metabolic network incorporated within each cell gives us the advantage of profiling each metabolite concentration with time within a discrete cell, as well within the population as a whole. We first looked at the spatially averaged behavior of each metabolite concentration to correlate with the behavior of the whole tumor. That is, we examined the total concentration of each metabolite for a slice along the width of the tumor. The metabolite concentration values from all 30 iterations were then averaged. We then plotted the averaged metabolite concentration along the slice, at successive time points.

We first examine the concentration profiles of the metabolites directly affected by the metabolic perturbations (Figure 6). For the control case, intra-tumoral glucose is high at early times, and then decreases after three days of tumor growth (Figure 6(a), Column I). Additionally, there is a slight rebound in the glucose concentration for the control case at day 15. For all of the knockdown strategies, the glucose concentration is also predicted to have a slight time-delayed decrease (Figure 6(a), all columns). However, the glucose level does not rebound when GAPDH, OXPHOS or ASCT2 are knocked down. In fact, with OXPHOS or ASCT2 knockdown, intracellular glucose goes to zero after approximately 12 days of growth.

**Fig. 6.**
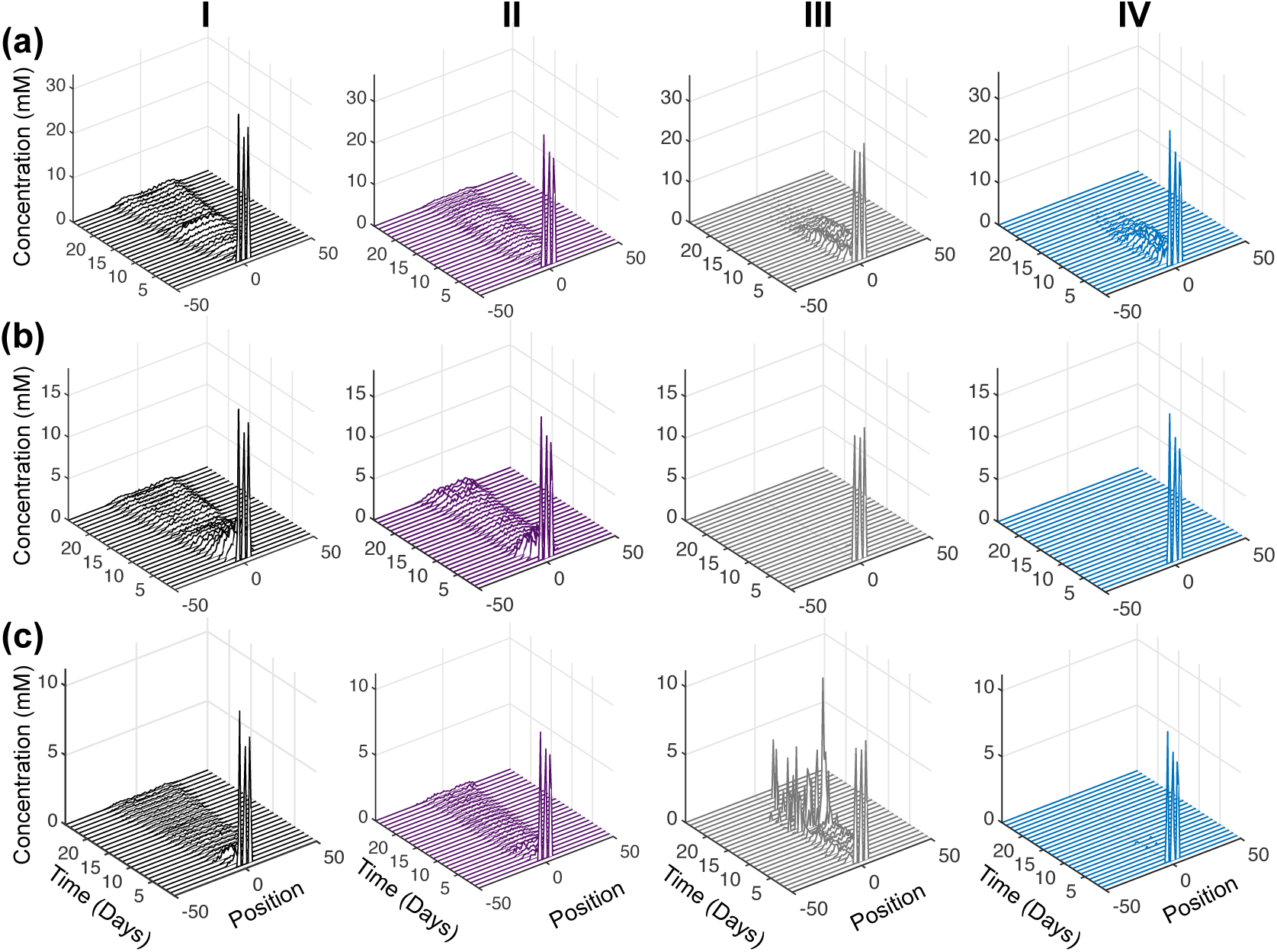
Intracellular Response of Complete Inhibition: Concentration profiles of total (a) Glucose, (b) ATP and (c) Glutamine in tumor slices along the *y*-axis in case of control with no inhibition (black, Column I), complete inhibition of GAPDH (purple, Column II), complete inhibition of OXPHOS (grey, Column III) and complete inhibition of ASCT2 (teal, Column IV). Each time profile is an average of 30 iterations of model simulation with the reaction velocity, *V_f_*, of the targeted reaction set equal to zero in each case.

The predicted intracellular ATP concentration also helps to explain the tumor growth profile. For the case of no inhibition, ATP is high at early times, and then decreases to a low level (Figure 6(a), Column I). Similar to the glucose concentration, ATP slightly increases after the perturbation, in this case, starting at day 10. When GAPDH is knocked down, the ATP concentration profile is similar to that of glucose (Figure 6(a), Column II). This non-zero concentration indicates there is an alternate production of ATP outside of the GADPH reaction. In contrast, knocking out OXPHOS or ASCT2 significantly affects the intracellular ATP level, which goes to zero in both cases (Figure 6(b), Columns III and IV).

An interesting response is observed in case of glutamine concentration, particularly for the metabolic perturbations. The glutamine concentration is similar to glucose and ATP for the control case, and intracellular glutamine is slightly reduced when GAPDH is knocked down (Figure 6(c), Columns I and II), compared to the control case. Interestingly, glutamine concentration is predicted to increase sharply when the OXPHOS reaction is inhibited (Figure 6(c), Column III), compensating for the loss of ATP (Figure 6(b), Column III). This shows that in initial phases of tumor growth, glutamine is used to provide metabolites for the TCA cycle, allowing the tumor reach a slightly larger volume when OXPHOS is knocked down, as compared to complete inhibition of ASCT2 (Figure 5). In contrast, intracellular glutamine levels are completely diminished when the ASCT2 reaction is inhibited (Figure 6(c), Column IV), leading to the lowest tumor volumes across the three metabolic perturbations (Figure 5).

An intriguing effect was noticed for the intracellular metabolite, fumarate, which has been identified as an oncometabolite (57). For the control case (no inhibition), fumarate concentration is high at early times and decreases over time, going to nearly zero between days 13 to 20 (Figure 7(a)). However, the concentration of fumarate then increases across the tumor until the simulation time ends. Although fumarate is not directly related to the targets of the three reactions that were inhibited (GAPDH, OXPHOS and ASCT2), the model predicts that its concentration is altered as a result of the metabolic inhibition. Complete knockdown of GAPDH has the effect of decreasing fumarate concentration throughout the tumor (Figure 7(b)), with the low level of fumarate (near zero) occurring more rapidly (at the 5th day) and for a longer period of time, compared to the control case. This reduction in the fumarate concentration correlates very well with the tumor growth profile, which is predicted to decrease after the 5th day and begin growing more rapidly after day 20. Inhibiting OXPHOS also has the effect of reducing the intra-cellular fumarate concentration throughout the tumor (Figure 7(c)). Interestingly, complete repression of ASCT2 also brings down the concentration levels of fumarate to nearly zero (Figure 7(d)), and its concentration remains very low, as compared to the control case. This matches the significantly lower tumor volume with ASCT2 knockdown compared to no inhibition. Overall, the time course of the fumarate concentration profile for each of the metabolic perturbations is linked to the changes in the overall growth of the tumor (Figure 5).

**Fig. 7.**
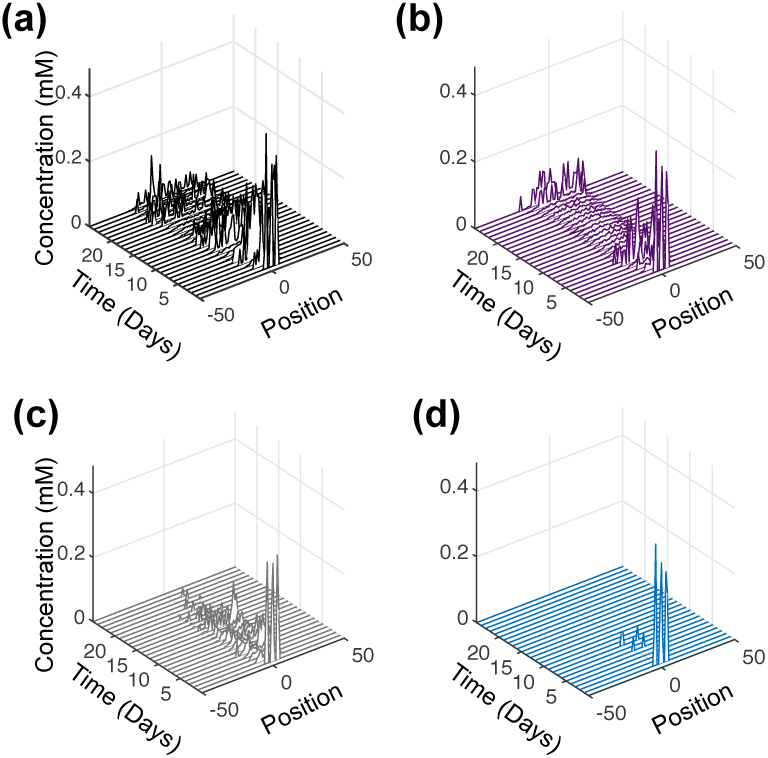
Intracellular Fumarate upon Complete Inhibition: Time profile of the total fumarate concentration in tumor slices along the *y*-axis. (a) Control with no inhibition (black), (b) complete inhibition of GAPDH (purple), (c) complete inhibition of OXPHOS (grey) and (d) complete inhibition of ASCT2 (teal). Each time profile is an average of 30 iterations of model simulations with the reaction velocity, *V_f_*, of the targeted reaction set equal to zero in each case.

#### Metabolic Perturbation II - Partial inhibition

The growth profile of the tumor was simulated to mimic time-based partial inhibition (90%) of three reactions - GAPDH, OXPHOS and ASCT2 (Figure 8). We again examined the intracellular concentration profiles of glucose, ATP and glutamine as potential explanations for the predicted tumor growth curves. To better understand the effect and obtain a visible correlation with the overall tumor growth, we plotted the time profile of glucose, ATP and glutamine aggregated over space, similar to the output from a homogenous population of cells (Figures 9 to 12).

**Fig. 8.**
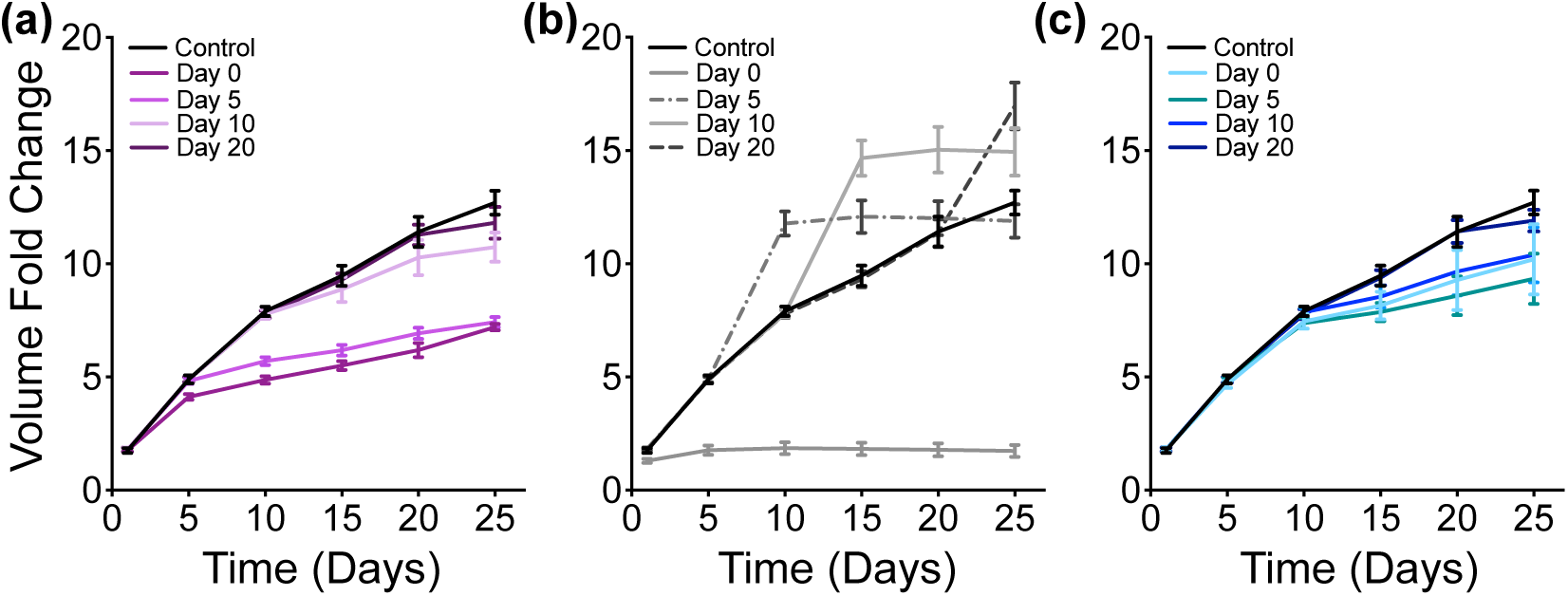
Metabolic Perturbation II - Partial Inhibition: Fold-change in tumor volume with time-based partial inhibition of (a) GAPDH (purple), (b) OXPHOS reaction (grey) and (c) ASCT2 (teal) compared to a control case (black). The error bars represent the standard deviation between 30 iterations of model simulations for each case.

**Fig. 9.**
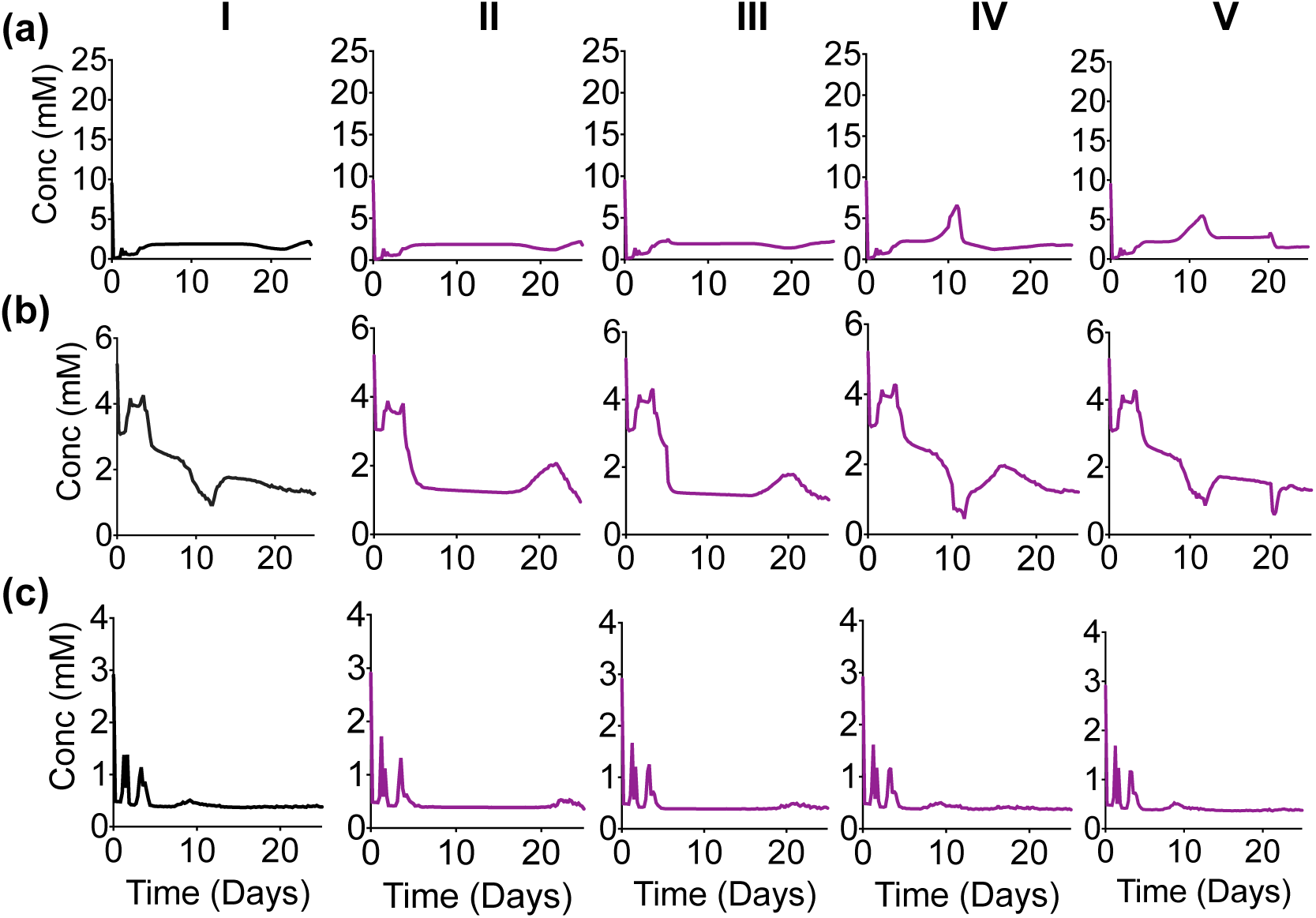
Intracellular Response of Partial Time-Based - GAPDH Inhibition: Spatially-aggregated concentration of intracellular metabolites: (a) Glucose, (b) ATP and (c) Glutamine. (Column I) No inhibition. With GAPDH inhibition at various times: (Column II) zeroth day, (Column III) 5th day, (Column IV) 10th day and (Column V) 20th day. Lines are an average of 30 iterations of the model simulation. For each iteration, the reaction velocity is taken from a Gaussian distribution based on the original velocity *V_f_*, with a mean value of 0.1 *\* V_f_* and a standard deviation of 0.2 *\* V_f_*.

#### GAPDH Inhibition

Partial repression of the GAPDH reaction at time zero (Day 0) shows a slightly delayed response in lowering the tumor volume. But when the reaction is inhibited at later time points (Day 5, 10 or 20), the effect of reducing the tumor volume is immediate (Figure 8(a)). Partial repression of GAPDH reaction at day zero in not seen to have any immediate effect on the concentration of intracellular glucose, since this species is upstream of the GAPDH reaction. Glucose concentration is predicted to increase after inhibition, as clearly seen when GAPDH is inhibited on the 10th day of growth (Figure 9(a), Column IV). In all cases of the time-based partial inhibition, the effect of targeting GADPH is only seen after the 5th day (Figure 9(a), Column II to V) compared to no inhibition (Figure 9(a), Column I), even if the inhibition is executed on day zero.

Intracellular ATP is minimally affected by the inhibition of GAPDH in the initial days (Figure 9(a), all Columns). For the control case, there is a decrease in ATP at day five and subsequent rebound in its concentration between days 10 and 15. When GAPDH inhibition occurs at Day 0 or 5, that rebound is delayed until after 15 days of growth (Figure 9(b), Columns II and III). Interestingly, if GAPDH is inhibited at Day 10 or 20, the rebound in the ATP concentration happens at nearly the same time as in the control case (Figure 9(b), Columns IV and V). This insight into the intracellular dynamics of key metabolites helps explain the overall tumor growth being similar to the no inhibition case (Figure 8(a)).

The concentration of glutamine is not noticeably affected by GAPDH inhibition, since it is far from the main target of this inhibition (Figure 9(c), all Columns). Thus, the tumor continues to produce the necessary metabolites to generate metabolites needed for tumor growth.

Overall, in the case of GAPDH inhibition, tumor volume initially follows the growth curve for the control case, and the effect of inhibiting the GAPDH reaction is only predicted to occur after approximately five days. This illustrates a growth surge during the initial phases of tumor growth, when the nutrients are present at sufficient levels. At this stage, a partial repression of GAPDH is not very successful in immediately producing the desired response of reduced tumor volume. However, executing the partial repression when the tumor has optimally utilized the nutrients (in our simulations, at Day 10 or 20) and has passed the growth surge is more effective in immediately reducing tumor volume (Figure 8(a)]. During the later stages of tumor growth even though the perturbations have an immediate effect on the intracellular concentrations of glucose and ATP, their concentrations seem to quickly relax back to the values they would have achieved if GAPDH were not inhibited at all. However, these effects on the intracellular metabolite concentrations are not pronounced enough to be reflected in the growth of the tumor as a whole, and tumor volume is only slightly decreased, compared to the control case.

These predictions provide mechanistic insight into the dynamic reprogramming that cells can exhibit at the intracellular level to respond and adjust to metabolic perturbations and continue along a similar growth trajectory as the control case.

#### OXPHOS Inhibition

The partial inhibition of OXPHOS on the zeroth day has an immediate effect on the tumor growth, which is significantly reduced compared to the control case (Figure 8(b)). However, the suppression of OXPHOS at times after the tumor has had a chance to grow has an antagonistic effect on tumor volume. Specifically, inhibiting the OXPHOS reaction at Day 5, 10 or 20 triggers an immediate increase in the tumor volume, after which the tumor volume remains steady.

We investigated the concentration profiles of key intracellular metabolites, spatially-averaged over the entire cell population (Figure 10), in order to explain this unexpected behavior. An interesting observation is seen for intracellular glucose. The slight increase in glucose concentration that was observed in case of time-delayed partial GAPDH repression is much more prominent for OXPHOS inhibition (Figure 10(a), Columns III to V). That is, the model predicts that targeting oxidative phosphorylation by reducing the reaction velocity of the OXPHOS reaction leads to increased glycolysis. It has been known that cancer cells have increased flux through the glycolysis pathway in the absence of oxygen (the *Pasteur effect* (51)), which is also observed even in the presence of oxygen (the *Warburg effect*). Thus, our simulations directly match experimental observations (16).

**Fig. 10.**
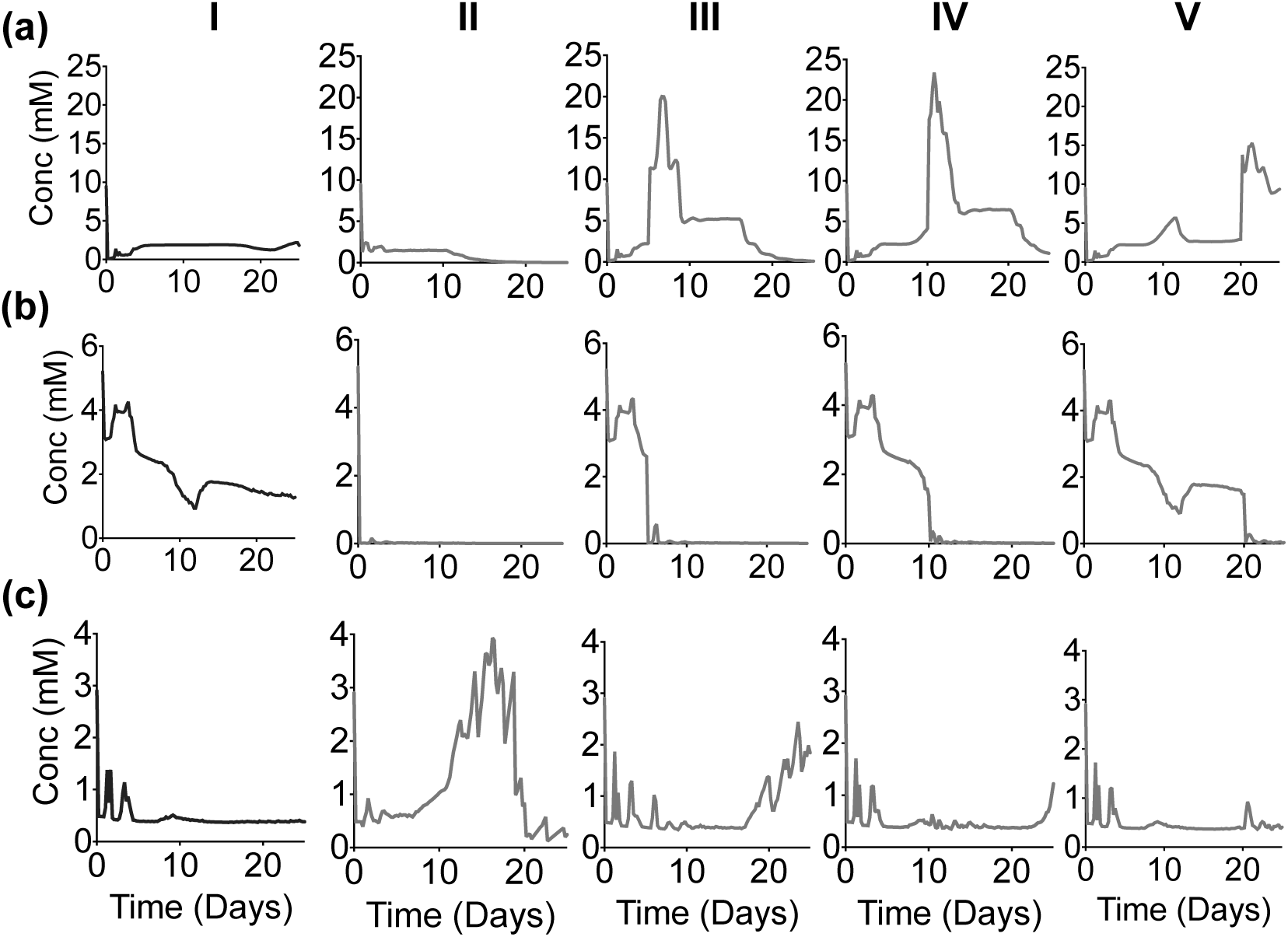
Intracellular Response of Partial Time-Based - OXPHOS Inhibition: Spatially-aggregated concentration of intracellular metabolites: (a) Glucose, (b) ATP and (c) Glutamine. (Column I) No inhibition. With OXPHOS inhibition at various times: (Column II) zeroth day, (Column III) 5th day, (Column IV) 10th day and (Column V) 20th day. Lines are an average of 30 iterations of the model simulation. For each iteration, the reaction velocity is taken from a Gaussian distribution based on the original velocity *V_f_*, with a mean value of 0.1 *\* V_f_* and a standard deviation of 0.2 *\* V_f_*.

The intracellular concentration of ATP immediately decreases once the OXPHOS reaction is inhibited, irrespective of the timing of the inhibition (Figure 10(b), all columns). However, the cells’ growth rate depends not only on the concentration of ATP, but also on intracellular glucose and glutamine, which can be used to produce cellular components for proliferating cells besides energy. Thus, tumor growth is still possible even in the absence of ATP.

Although OXPHOS inhibition has an immediate effect on the intracellular concentrations of glucose and ATP (Figure 10(a)-(b)), any noticeable effects on glutamine concentration are more delayed (Figure 10(c)). The concentration of intracellular glutamine is predicted to increase when OXPHOS is inhibited on the zeroth day (Figure 10(c), Column II). For this case, glutamine concentration increases after both glucose and ATP are depleted; however, the tumor volume remains low (Figure 8(b)). This indicates that increased glutamine is not sufficient to enable tumor growth. Rather, glutamine is used to maintain the tumor volume. This also occurs for the time-delayed inhibition. For example, when OXPHOS is inhibited at Day 5 (Figure 10(c), Column III), the increased glucose concentration leads to an increase in the tumor growth rate (between days 5 to 10). But once glucose is depleted at day 18, the tumor volume remains constant, even though glutamine concentration begins to increase at the same time.

In addition to investigating the concentrations of the key metabolites used for proliferation, We also look to glycolytic intermediates to explain the increased tumor growth after OXPHOS inhibition (Figure 11). The increased reliance on glycolysis when the OXPHOS reaction is inhibited (indicated by the higher glucose levels shown in Figure 10(a)) is also reflected in the concentrations of PEP (Figure 11(a)) and lactate (Figure 11(b)). Specifically, PEP and lactate are both increased when OXPHOS is inhibited, complementing the glucose concentration profile. The predicted increases in PEP and lactate are consistent with experimental observations (58). Altogether, the simulation results show that with OXPHOS inhibition, the cells become more glycolytic as an alternative mechanism to produce the cellular components needed for tumor growth.

**Fig. 11.**
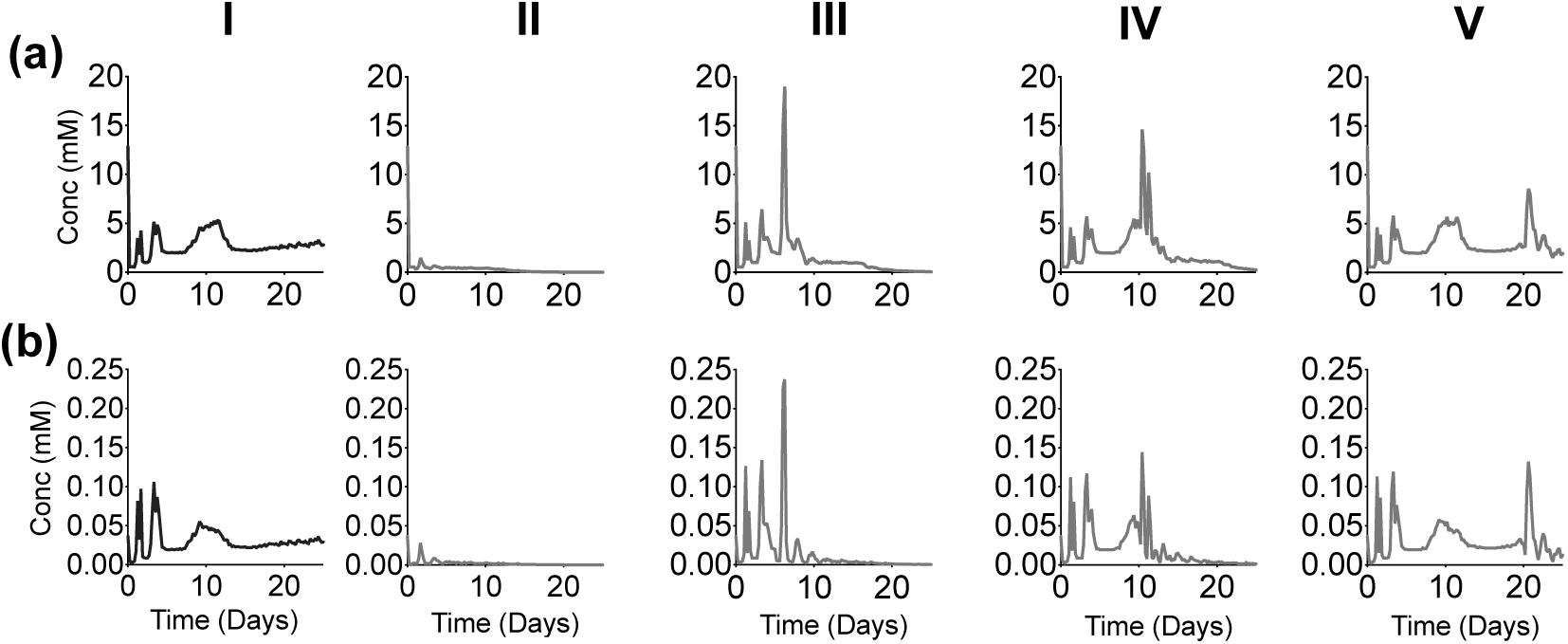
Up-regulation of glycolytic pathway: Spatially-aggregated concentration of intracellular metabolites: (a) PEP and (b) Lactate (Column I) No inhibition or with GAPDH inhibition at various times: (Column II) zeroth day, (Column III) 5th day, (Column IV) 10th day and (Column V) 20th day.

In summary, by studying the intracellular concentration of glucose, ATP and glutamine, we find that simulating inhibition of the OXPHOS reaction demonstrates the ability of the cancer cells to rewire their pathways to sustain tumor growth. Specifically, the simulations show that the primary fuel for cell proliferation and overall tumor growth switches from ATP to glucose to glutamine, all in a relatively short timescale (on the order of hours to days).

#### ASCT2 Inhibition

Partial inhibition of the ASCT2 reaction does not have an immediate effect on tumor volume (Figure 8(c)). This indicates the preferential dependence of the tumor cells on glycolysis and mitochondrial production of ATP during its initial phase. It also points towards the cell’s ability to utilize the minimal glutamine available due to the remaining 10% activation of the ASCT2 reaction to provide the necessary nutrient for cell growth.

By investigating the intracellular metabolite concentrations, the model predicts that glucose is not affected by the inhibition of ASCT2 (Figure 12(a)). In addition, the ATP concentration is robust to ASCT2 inhibition, as the ATP concentration profile is nearly the same across all four cases (Figure 12(b)). Only when the ASCT2 reaction is inhibited at Day 10 or 20 (Figure 12(b), Columns IV and V) does the ATP concentration exhibit a sharper decrease from Days 10 to 12, compared to inhibition at Day 0 or 5 (Figure 12(b), Columns II and III). Interestingly, the partial inhibition of ASCT2 does not have a noticeable direct effect on the concentration of glutamine (Figure 12(c)), further emphasizing that fact that even a 10% activation of ASCT2 pathway is enough to sustain enough glutamine supply for the downstream pathways to remain functional.

**Fig. 12.**
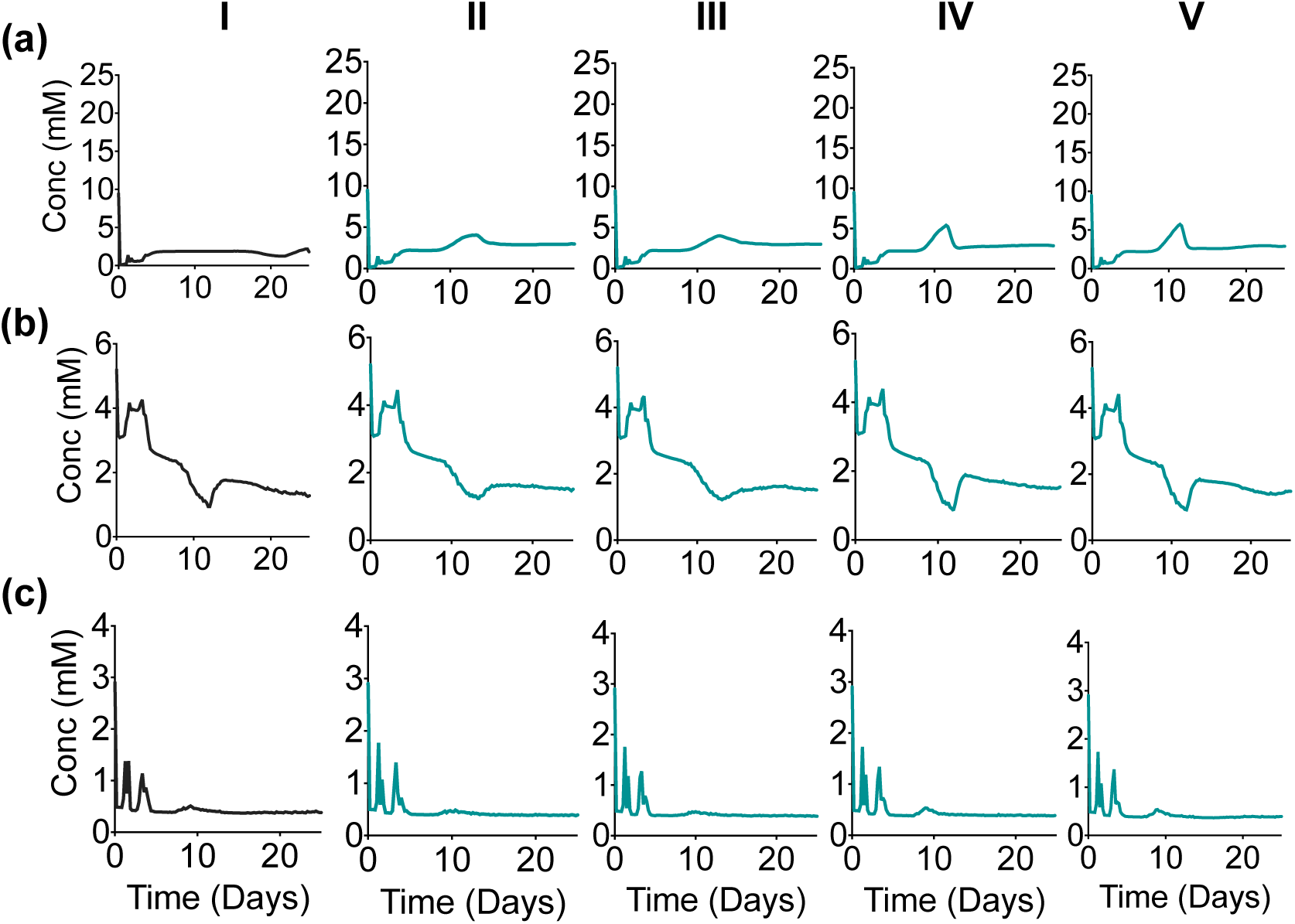
Intracellular Response of Partial Time-Based - ASCT2 Inhibition: Spatially-aggregated concentration of intracellular metabolites: (a) Glucose, (b) ATP and (c) Glutamine. (Column I) No inhibition. With ASCT2 inhibition at various times: (Column II) zeroth day, (Column III) 5th day, (Column IV) 10th day and (Column V) 20th day. Lines are an average of 30 iterations of the model simulation. For each iteration, the reaction velocity is taken from a Gaussian distribution based on the original velocity *V_f_*, with a mean value of 0.1 *\* V_f_* and a standard deviation of 0.2 *\* V_f_*.

## Discussion

In this paper, we present a multiscale model of tumor growth and use the model to investigate the effect of targeted perturbations of intracellular metabolism on the growth of the tumor as a whole. We simulate tumor growth initiating from a discrete cluster of cells. Our cell-based model incorporates three levels of cellular decision-making processes integrated in a multiscale manner. At the extracellular level, a system of partial differential equations models nutrient diffusion. At the cellular level, a discrete Cellular Potts modeling approach is used to simulate cellular motility and inter-cellular interactions. At the sub-cellular level, a detailed kinetic model in the form of ordinary differential equations accounts for intracellular metabolism. Together, these processes contribute to the initial cluster of tumor cells increasing in number and volume. Specifically, the cells take up extracellular nutrients and use them to fuel proliferation via the metabolic network. In our simulations, a small cluster of tumor cells evolves into a layered tumor consisting of proliferating and quiescent cells surrounding a central core of necrotic cells. At any point in time, the tumor volume and total number of viable cells depend not only on the surrounding extracellular environment but also the response of each discrete cell due to its intracellular state.

Our model is able to reproduce the growth profile of an avascular tumor and identify a robust set of parameters. Due to the computational expense in simulating a detailed mechanistic model of the metabolic network within each discrete cell, we chose not to model tumor vascularization. In the avascular stage of tumor growth, the tumor depends on the nutrient supply from neighboring tissues. We achieve this in our model by letting the stromal tissue layer (ECM and Basal cells) secrete glucose, glutamine and oxygen. During the initial stage of growth, an avascular tumor exhibits “quasi-exponential” growth followed by a saturation phase in which it maintains a constant volume, since without angiogenesis, these avascular tumors are not capable of procuring enough nutrients to continue an exponential growth (59). Our model of avascular tumor growth produces spheroid volumes and growth profiles that are in agreement with experimental data, where the set of parameters needed to match the data was identified by sampling the space for parameters that significantly influence the tumor growth. Given this confidence in the model and the identified parameter sets, we applied the model to investigate various aspects of tumor growth.

We find that tumor growth is more dependent on the intracellular ATP threshold values than on lactate, emphasizing that ATP is more important in the initial growth of tumor. Lowering the threshold values of ATP at which a quiescent cell can transition into a proliferating cell that continues to grow and divide by 10-fold, increases the tumor volume by almost three-fold. Interestingly, the heterogeneity in tumor growth profile at low and high values of the ATP threshold is more pronounced, as a higher standard deviation is observed in tumor volumes during iterative simulation runs.

We next applied the model to investigate the effects of phenotypic heterogeneity in tumors originating from slight variations in the intracellular conditions within each discrete cell. The model simulations show that the variation in intracellular conditions do not produce a noticeable impact on tumor volume during the initial phase of avascular tumor growth. However, the effects of different intracellular initial conditions for each cell are more prominent during the later stages of growth. Specifically, there was a high standard deviation for the average tumor growth profile, indicating the divergence of individual growth trajectories.

Most excitingly, we applied the model to understand the complex intracellular responses following targeted metabolic perturbations. That is, we simulated the effects of inhibiting the uptake of major nutrients (glucose and glutamine) as well as the primary reaction used for ATP production. Since each cell consists of a detailed mechanistic metabolic network, the model can be used to simulate inhibition of the specific target within each discrete cell and to predict how those perturbations affect the behavior of the tumor as a whole. We simulated targeted inhibition of: (a) the GAPDH reaction, a glycolytic pathway reaction responsible for converting glyceraldehyde 3-phosphate to 1,3-bisphosphoglyceric acid, (b) the OXPHOS reaction, which is responsible for the conversion of ADP to ATP, representing mitochondrial ATP production and (c) the ASCT2 reaction, an enzymatic reaction responsible for the uptake of extracellular glutamine. While a complete repression of these reactions had an expected outcome of reducing tumor growth, the model simulations generated an unexpected response for the tumor volume when partial time-based inhibition of OXPHOS reaction was implemented. Specifically, inhibiting OXPHOS at various times after tumor growth had begun, enables the cancer cells to dynamically change their reliance on the available nutrients and continue growing (“metabolic reprogramming”). Through the simulated perturbations, the model provides direct insight into how the cells up-regulate glucose utilization via the glycolytic reactions when deprived of ATP (as reflected in the increased concentrations of PEP and lactate). Interestingly, this increased glycolysis does not take place if the cells are deprived of ATP at the start of tumor growth. Rather, the reprogramming becomes apparent when the ATP deprivation occurs during the exponential growth phase.

In addition, our model simulations of various metabolic perturbations provided a first-hand proof of preferentially active pathways during tumor progression. As the tumor initiates its growth, every cell has sufficient nutrient availability. This is supported by the complete lowering of tumor growth when oxidative phosphorylation or glutamine uptake is completely inhibited. However, the tumor growth is not as affected with complete repression of GAPDH. We suspect that this is because of two reasons. First, GAPDH is downstream in the glycolytic pathway, and a complete repression of a downstream target takes time to be effective. Second, we hypothesize that glycolysis becomes a major player to support tumor growth at intermediate phases, most importantly, during hypoxia (as modeled by inhibition of OXPHOS). The glycolytic pathway is up-regulated under partial repression of oxidative phosphorylation during the exponential growth phase. During this phase, the tumor alters its metabolic pathway to compensate for the loss of one nutrient by maximizing an alternative pathway to supply ATP.

Additionally, our model simulations demonstrate clearly the role of glutamine in tumor growth as an alternative but necessary nutrient. Glutaminolysis provides the cells with carbons to replenish TCA cycle intermediates and generate building blocks for cell growth. While complete inhibition of the glutamine uptake (the ASCT2 reaction) has a profound influence on tumor growth, partial inhibition of glutamine uptake prompts the cells to utilize the minimal glutamine available, as well as rely more heavily on glucose and generation of ATP to sustain tumor growth. Thus, the model predictions indicate that during avascular growth, the tumor cells heavily depend on oxidative phosphorylation and glutaminolysis.

Taken together, our simulations for the effects of metabolic perturbations indicate that while it is important to identify the optimal targets for inhibiting tumor growth, it is equally necessary to identify the time at which the targeted therapies should be administered. Although a single optimal target inhibition may be sufficient to impede tumor growth during the initial phases of tumor growth, that same target may not be as effective during later stages of tumor growth. At this stage, a synergistic combinatorial therapies targeting various pathways of nutrient utilization and ATP production could be more successful to impede tumor growth.

We acknowledge some limitations of our model. Firstly, due to the high computational expense in simulating a detailed sub-cellular kinetic network in a large number of cells, vascular growth (angiogenesis) was not modeled. However, given the multiscale nature of our modeling approach, this limitation may be addressed in future extension of the model by incorporating vascularization or angiogenesis through a signaling network at a different timescale. Secondly, we can expand the model to more explicitly account for “heterocellular interactions” (60) that occur between different cells. This means including interactions between stromal and malignant cells, not only in terms of the availability of nutrients but also how these close interactions influence the metabolism dynamics within each cell. We currently have a static metabolic network within each cell, but that network can be rewired and influenced by neighboring cells. Finally, our current model does not simulate metastasis. We provide the basic framework that could be further extended to include metastasis and invasion into the epithelial layer by enabling the daughter cells to have a random distribution of growth rates and adhesivity values, increasing stochasticity and heterogeneity and providing a more realistic picture of pancreatic tumor growth.

## Conclusion

Altered cellular metabolism is a hallmark of tumorigenicity and malignancy. Mathematical models serve as “non-invasive” tools (61) to understand better the prevalent heterogeneities in tumors and assess the potential effects of various therapeutic strategies. We developed a multiscale cellular model of avascular tumor growth and apply it to predict the effects of how targeting specific intracellular metabolic reactions influences tumor growth. The particularly novel aspect of our model is the detailed metabolic network at the sub-cellular level that directly influences growth at the cellular and tissue levels. The multiscale cell-based model presented in this study enables an investigation of how cellular-level features impact the evolution of a tumor. In addition, the stochastic modeling approach provides a more detailed picture of both intrinsic and extrinsic heterogeneities that are prevalent in tumors. We modeled the response of the tumor as a whole to several targeted perturbations and provide mechanistic explanations of the response as a function of the intra-cellular dynamics of metabolism. Our model is a useful tool to screen potential therapeutic strategies in cancer, complementing experimental studies.

## Supporting information

## ACKNOWLEDGEMENTS

This work is funded by The Rose Hills Foundation and the USC Provost’s Office (research grant to S.D.F.). Simulations were performed at the Center for High-Performance Computing of the University of Southern California.

## Supporting Information

**S1 Text. Supplementary Material.** Supplemental Tables and Figures, along with detailed explanation of parameters listed in Table 1.

**S1 Movie. Representative movie of tumor growth.**

