## Supplementary material for "Metabolic reprogramming dynamics in tumor spheroids: Insights from a multicellular, multiscale model"

### Supplementary Tables

**Table S1. Values of constants used in the model simulations**

| Quantity | Significance | Model values |
| --- | --- | --- |
| 1 pixel | Unit length of grid | 4 $\mu$m |
| 1 voxel | Unit surface area of grid | 16 $\mu m^2$ |
| 1 voxel | Unit volume of grid | 64 $\mu m^3$ |
| NeighborOrder | Interaction Range | 3 |
| $T_m$ | Internal fluctuation amplitude | 50 |
| $\alpha$ | Secretion rate of nutrients | 0.5 fmol/voxel (0.0222 fmol/cell/sec) |
| Glucose in blood | Steady State Concentration | 5 mM (0.32 fmol/voxel) |
| Oxygen in blood | Steady State Concentration | 0.056 mM (0.003584 fmol/voxel) |
| Glutamine in blood | Steady State Concentration | 1 mM (0.064 fmol/voxel) |
| Lactate in blood | Steady State Concentration | 2 mM (0.128 fmol/voxel) |
| $\epsilon_{Glc}$ | Decay rate of glucose | 1.562/mcs |
| $\epsilon_{Oxy}$ | Decay rate of oxygen | 139.5089/mcs (1) |
| $\epsilon_{Gln}$ | Decay rate of glutamine | 7.8125/mcs (2) |
| $\epsilon_{Lac}$ | Decay rate of lactate | 3.9062/mcs (3) |
| $D_{glc}$ | Diffusion constant of glucose | 500 $\mu m^2$/s (11250 voxel/mcs)) |
| $D_{oxy}$ | Diffusion constant of oxygen | 1820 $\mu m^2$/s (40950 voxel/mcs)) (1) |
| $D_{gln}$ | Diffusion constant of glutamine | 567 $\mu m^2$/s (12575.5 voxel/mcs)) |
| $D_{lac}$ | Diffusion constant of lactate | 178 $\mu m^2$/s (4005 voxel/mcs)) (4) |
| $\lambda$ | Chemotaxis coefficient | 500 |

**Table S2. Fixed set of parameter values**

| Parameters | Significance | Values |
|---|---|---|
| V0 | Target Volume | 16 |
| S0 | Target Surface | 16 |
| $\lambda_{V0}$ | Volume Constraint | 15 |
| $\lambda_{S0}$ | Surface Constraint | 5 |
| $V0_{nec}$ | Target Volume of a necrotic cell | 4 |
| $S0_{nec}$ | Target Surface of a necrotic cell | 4 |
| $\lambda_{NV0}$ | Volume Constraint of necrotic cell | 10 |
| $\lambda_{NS0}$ | Surface Constraint of necrotic cell | 5 |
| EV0 | Target Volume of an epithelial cell | 40*V0 |
| ES0 | Target Surface of an epithelial cell | 40*S0 |
| $\lambda_{EV0}$ | Volume Constraint of an epithelial cell | 40*$\lambda_{V0}$ |
| $\lambda_{ES0}$ | Surface Constraint of an epithelial cell | 40*$\lambda_{S0}$ |
| kgts | surface = kgts*sqrt(volume) | 4 |
| Pvolmaxmit | Maximum volume of a proliferating cell | 2*V0 |
| Svolmaxmit | Maximum surface of a proliferating cell | 2*S0 |
| MaxDistance | Distance between COM of two epithelial cells | 4.5 |
| TargetDistance | Target Distance between two epithelial cells | 4.0 |
| $\lambda_{ecm-ecm}$ | Distance constraint between ECM cells | 60 |
| $\lambda_{ecm-basal}$ | Distance constraint between ECM and Basal cells | 40 |
| $\lambda_{basal-basal}$ | Distance constraint between Basal cells | 60 |
| QCNeThr | QCancer to Necrotic transition threshold | 2*PNeThr |
| PSNeThr | PStem to Necrotic transition threshold | 4*PNeThr |
| QSNeThr | QStem to Necrotic transition threshold | 8*PNeThr |

**Table S3. Adhesion Coefficients for different cell types**

| Cell Types | Medium | PCancer | QCancer | PStem | QStem | Necrotic | Basal | ECM |
|---|---|---|---|---|---|---|---|---|
| **Medium** | 0 | 10 | 10 | 10 | 10 | 10 | 2 | 2 |
| **PCancer** | 10 | 2 | 10 | 10 | 10 | 10 | 6 | 6 |
| **QCancer** | 10 | 10 | 2 | 10 | 10 | 10 | 8 | 8 |
| **PStem** | 10 | 10 | 10 | 2 | 10 | 10 | 6 | 6 |
| **QStem** | 10 | 10 | 10 | 10 | 2 | 10 | 8 | 8 |
| **Necrotic** | 20 | 10 | 10 | 10 | 10 | 2 | 20 | 20 |
| **Basal** | 2 | 6 | 8 | 6 | 8 | 20 | 10 | 16 |
| **ECM** | 2 | 6 | 8 | 6 | 8 | 20 | 16 | 10 |

### Supplementary Figures

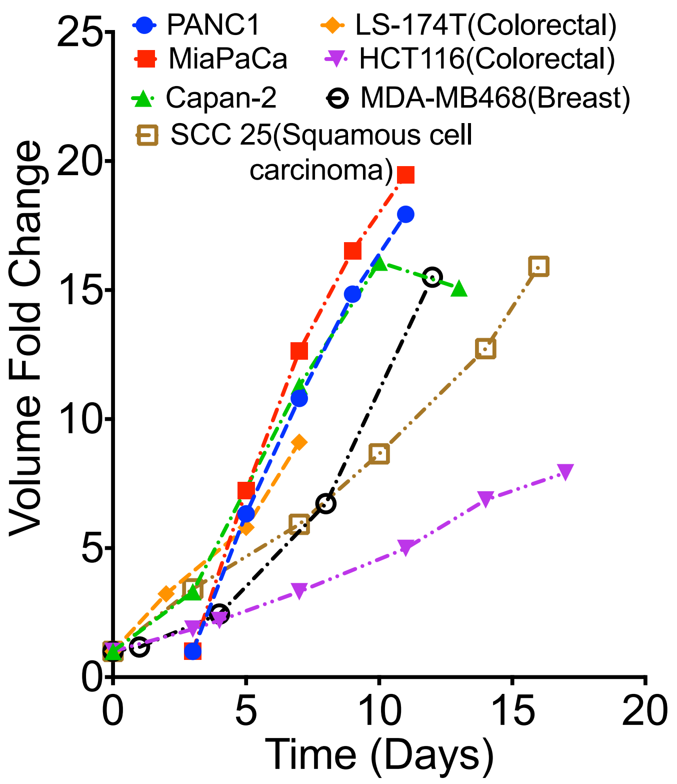


**Figure S1. Experimental data of *in vitro* tumor spheroids for various cancer cell lines.** With a range of growth profiles (5–7) observed for different multicellular tumor spheroids generated from human cancer cell lines under various growth conditions, a 10-fold change in volume in approximately 15 days was set as the standard for calibrating the model simulations to experimental measurements.

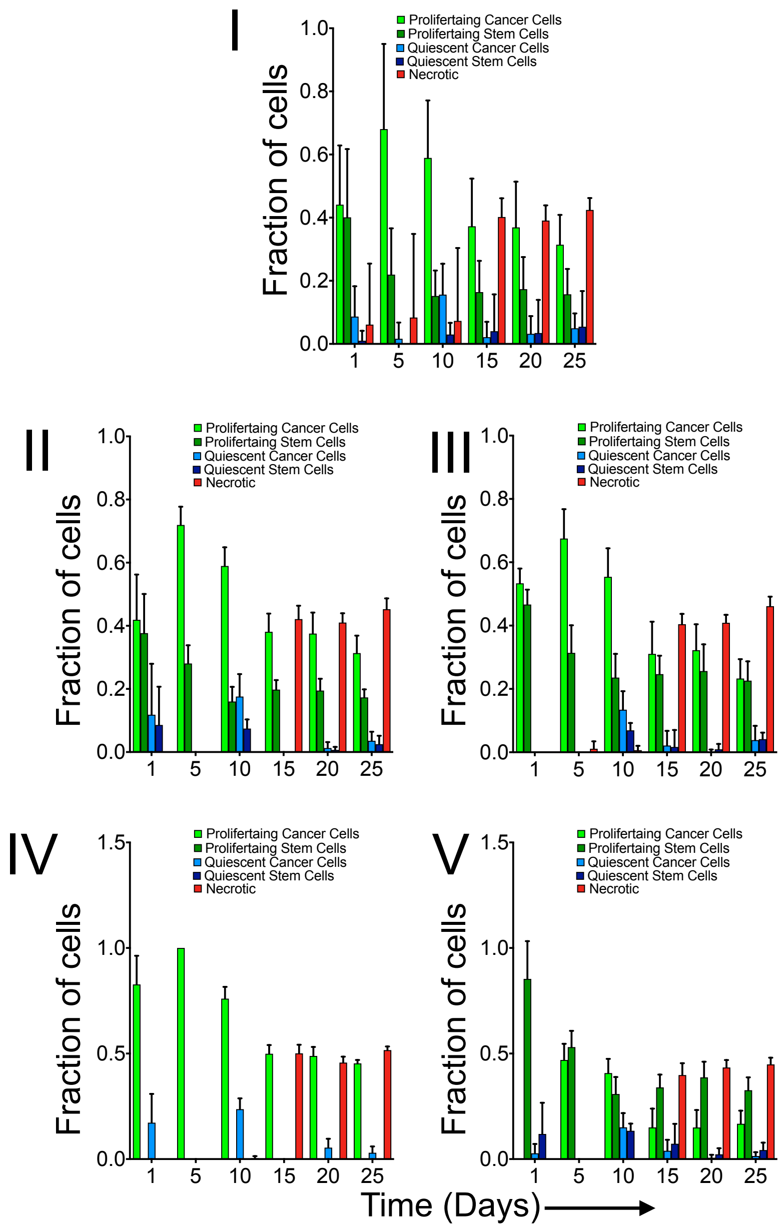


**Figure S2. Distribution of different cell types.** The number of each cell type varies with time when the tumor is initiated with different configurations (see Figure 3(a)): (I) All four types of cells (PCancer, QCancer, PStem and QStem), (II) Only proliferating cell (PCancer and PStem), (III) Only quiescent cells (QCancer and QStem), (IV) Only proliferating cancer cell (PCancer) and (V) Only proliferating stem cells (PStem).

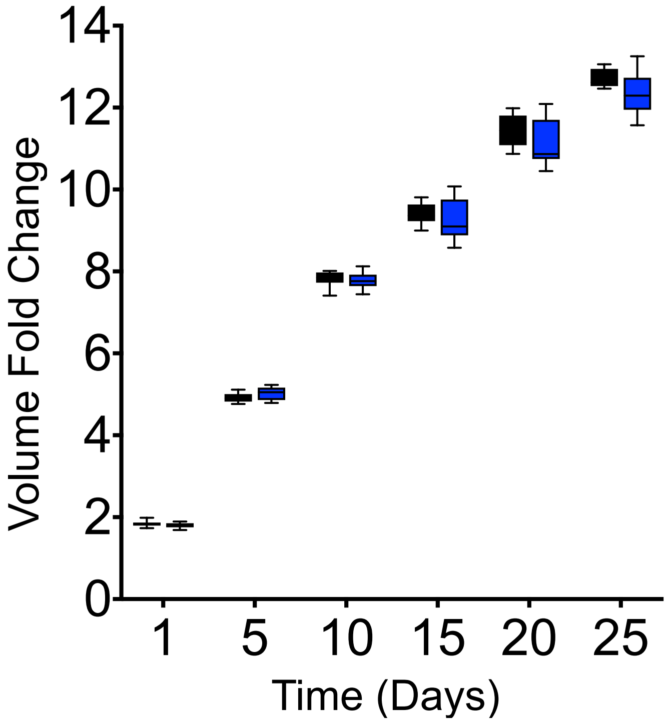


**Figure S3. Simulating different initial metabolite concentrations.** Model simulations obtained when each cell is initiated with a slight variation in the intracellular conditions, eventually generating greater inter-tumor heterogeneity as the tumor grows. The figure depicts the time profile of tumor growth for 10 iterations of the control case where each simulation starts with same intracellular metabolite concentrations for each cell in the initial cluster (black) and 10 iterations where each cell has a different set of initial conditions (blue). The error bars represent the standard deviation.

### Description of parameters given in Table 1 of the main text

1. *incvol*: The rate at which cells increase their volume to proliferate, a property of only proliferating cells (PCancer and PStem). The higher the rate, the greater the increase in the cell's volume, leading to a higher rate of proliferation.

2. *decvol*: The rate at which necrotic cells decrease their volume, mimicking apoptosis.

3. *PGrThr*: The threshold value for the total concentration of ATP, intracellular glucose and glutamine needed for a proliferating cell to increase its volume.

4. *SGrThr*: The threshold value for the total concentration of ATP, intracellular glucose and glutamine needed for a stem cell to increase its volume.

5. *StressIncrement*: The value by which the Stress attribute is increased when the required number of neighbors (*N*) surround a cell and exert compressive stress.

6. *StressThr*: The threshold value of the Stress attribute that the cells have to cross to transition into a necrotic cell.

7. *N*: The number of neighbors that a cell has, which influences the accumulation of Stress.

8. $V_{atpmax}$: A parameter influencing how cell attributes depend on ATP.

9. *atpD*: Threshold value below which the Starvation attribute is incremented for the cell, leading to necrosis and above which Health is acquired, leading to the conversion of a quiescent cell to a proliferating cell.

10. $PNeThr$: This is the threshold for a proliferating cell to transition into necrotic state due to a lack of ATP.

11. $PNeThr_{acidosis}$: This is the threshold for a proliferating cell to transition into necrotic state due to excessive exposure to an acidic environment.

12. $Total_{time}$: The total time for which the cells have to be exposed to a low ($Neg_{concATP}$) or high ($Pos_{concATP}$) concentration of ATP in order to transition to a necrotic or proliferating cell.

13. $Neg_{concATP}$: The low concentration of ATP used in the calculation of *PNeThr*. A low value of $Neg_{concATP}$ leads to low value of *PNeThr* and an easier transition of a proliferating cell to a necrotic cell. Alternatively, a high value of $Neg_{concATP}$ leads to high value of *PNeThr* respectively and an harder transition of a proliferating cell to a necrotic cell.

14. $Pos_{concATP}$: Total amount of intracellular ATP, glucose and glutamine used in the calculation of *QCPThr* and *QSSThr*, which affects the transition from a quiescent cell to a proliferating cell.

15. $V_{lacmax}$: A parameter for attribute factor calculation dependent on lactate.

16. *LacDeath*: Threshold value of lactate concentration, above which the Starvation attribute is incremented for the cell, leading to necrosis.

17. $Pos_{concLac}$: The high concentration of lactate used in the calculation of $PNeThr_{acidosis}$.

18. $Total_{timelac}$: This is the total time for which the cells have to exposed to high ($Pos_{concLac}$) concentration of lactate to transition to necrotic state due to acidosis.

19. *maxdiv*: Maximum number of divisions the cell can have before senescence sets in and the cell transitions to a necrotic cell.

20. *probstem*: The probability of a proliferating stem cell (PStem) to divide into a quiescent cancer (QCancer) or quiescent stem (QStem) cell.

21. *C*: This measures the amount of contribution of nutrients towards the accumulation of Health. Higher *C* values leads to faster accumulation of Health and an easier transition from a quiescent cell to a proliferating cell.

22. *a*: The Hill coefficient representing the cooperativity between nutrients responsible for the growth and transition of a cell.
