## Supplementary figures and images for "Metabolic reprogramming dynamics in tumor spheroids: Insights from a multicellular, multiscale model"

### Supplementary file 2

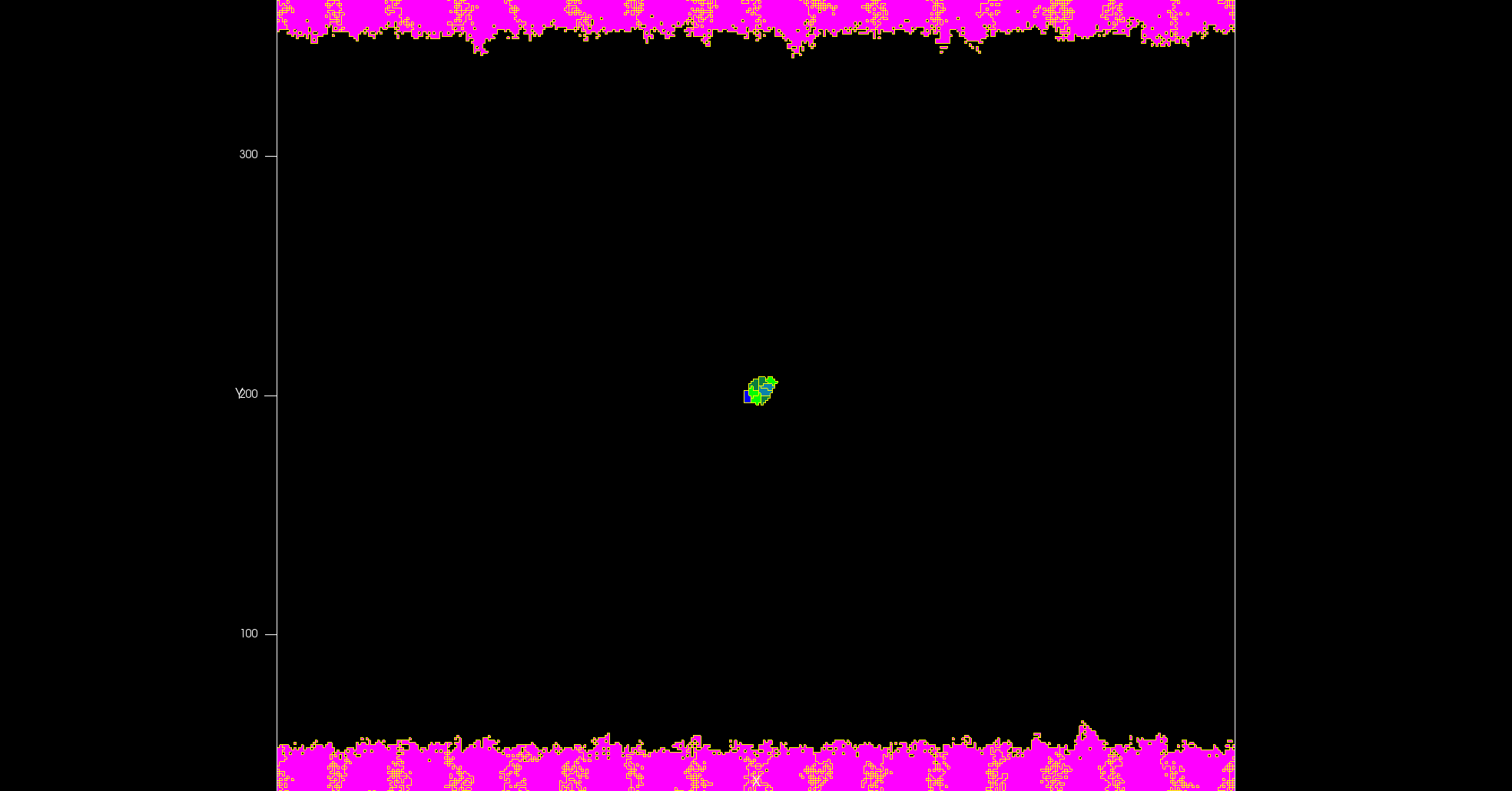
